# “Patient-specific Alzheimer-like pathology in trisomy 21 cerebral organoids reveals BACE2 as a gene-dose-sensitive AD-suppressor in human brain”

**DOI:** 10.1101/2020.01.29.918037

**Authors:** Ivan Alić, Pollyanna A Goh, Aoife Murray, Erik Portelius, Eleni Gkanatsiou, Gillian Gough, Kin Y Mok, David Koschut, Reinhard Brunmeir, Yee Jie Yeap, Niamh L O’Brien, Jurgen Groet, Xiaowei Shao, Steven Havlicek, N Ray Dunn, Hlin Kvartsberg, Gunnar Brinkmalm, Rosalyn Hithersay, Carla Startin, Sarah Hamburg, Margaret Phillips, Konstantin Pervushin, Mark Turmaine, David Wallon, Anne Rovelet-Lecrux, Hilkka Soininen, Emanuela Volpi, Joanne E Martin, Jia Nee Foo, David L Becker, Agueda Rostagno, Jorge Ghiso, Željka Krsnik, Goran Šimić, Ivica Kostović, Dinko Mitrečić, LonDownS Consortium, Paul T Francis, Kaj Blennow, Andre Strydom, John Hardy, Henrik Zetterberg, Dean Nižetić

## Abstract

A population of >6 million people worldwide at high risk of Alzheimer’s disease (AD) are those with Down Syndrome (DS, caused by trisomy 21 (T21)), 70% of whom develop dementia during lifetime, caused by an extra copy of β-amyloid-(Aβ)-precursor-protein gene. We report AD-like pathology in cerebral organoids grown *in vitro* from non-invasively sampled strands of hair from 71% of DS donors. The pathology consisted of extracellular diffuse and fibrillar Aβ deposits, hyperphosphorylated/pathologically conformed Tau, and premature neuronal loss.

Presence/absence of AD-like pathology was donor-specific (reproducible between individual organoids/iPSC lines/experiments). Pathology could be triggered in pathology-negative T21 organoids by CRISPR/Cas9-mediated elimination of the third copy of chromosome-21-gene *BACE2*, but prevented by combined chemical β and γ-secretase inhibition. We found that T21-organoids secrete increased proportions of Aβ-preventing (Aβ1-19) and Aβ-degradation products (Aβ1-20 and Aβ1-34). We show these profiles mirror in cerebrospinal fluid of people with DS. We demonstrate that this protective mechanism is mediated by BACE2-trisomy and cross-inhibited by clinically trialled BACE1-inhibitors. Combined, our data prove the physiological role of *BACE2* as a dose-sensitive AD-suppressor gene, potentially explaining the dementia delay in ∼30% of people with DS. We also show that DS cerebral organoids could be explored as pre-morbid AD-risk population detector and a system for hypothesis-free drug screens as well as identification of natural suppressor genes for neurodegenerative diseases.

## Introduction

Production^1–3^, and degradation^4^ of β-amyloid peptides (Aβ) are among the central processes in the pathogenesis of Alzheimer’s disease (AD). The canonical Aβ peptide is produced after sequential cleavage of the β-amyloid precursor protein (APP) by β-secretase and γ-secretase, generating a peptide that most often begins 99 amino acids (aa) from the C-terminus of APP with Asp1 and contains the next 37-42 aa of the APP sequence, generating a range of peptides (Aβ1-37, 38, 39, 40 and 42). The longer of these peptides can be detected in toxic amyloid aggregates in the brain, associated with AD and other neurodegenerative disorders^5^. As APP gene is located on human chromosome 21, people with Down Syndrome (DS, caused by trisomy 21 (T21)) are born with one extra copy of this gene, which increases their risk of developing AD. Non-DS (euploid) people inheriting triplication of the *APP* gene alone (Dup*APP*) develop AD symptoms by age 60 with 100% penetrance. Paradoxically, only ∼70% of people with DS develop clinical dementia by age 60, suggesting the presence of other unknown chromosome 21-located genes that modulate the age of dementia onset^6, 7^. A number of secretases participate in the physiological cleavage of APP^1, 8^, generating various peptides involved in neuronal pathology.

BACE1 is the main β-secretase in the brain^9^, while the expression and function of its homologue BACE2 (encoded by a chromosome 21 gene) remain less clear^10, 11^. At least 3 different activities of BACE2 were recorded with regards to APP processing: as an auxiliary β-secretase (pro-amyloidogenic), as a θ-secretase (degrading the β-CTF and preventing the formation of Aβ), and as Aβ-degrading protease (AβDP) (degrading synthetic Aβ-peptides at extremely acidic pH). It remains unclear which of these activities reflect the role of BACE2 in AD. The potential activity of BACE2 as an anti-amyloidogenic θ-secretase can be predicted from studies on a variety of transfected cell lines that overexpress *APP*, and artificially manipulate the dose of BACE2^12–15^.

These studies uncovered that BACE2 can cleave the product of β-secretase (APP β-CTF) between aa19 and aa20, generating a 1-19 fragment^13–15^, thereby potentially preventing the formation of amyloidogenic Aβ, and degrading the β-CTF that has been implicated in neuronal toxicity, and impairment of several neuronal functions, such as axonal transport and autophagy^16^. When offered synthetic Aβ40/42 peptides in solution, purified BACE2 protein can rapidly degrade them by cutting after aa20 and aa34, to generate the 1-20 and 1-34 peptide products, but only at very acidic pH (3.5-4). In this reaction, BACE2 is 150-fold more efficient than BACE1, which is also capable of this cleavage, upon conditions of increased enzyme concentration/time^12, 14^. Neither of these two putative anti-amyloidogenic actions of BACE2 (the θ-secretase activity, generating aa1-19, or the Aβ-degrading-protease activity (AβDP or Aβ clearance) generating aa1-20 and aa1-34), have yet been demonstrated to be the functional role of BACE2 under physiologically fluctuating gene doses *in vivo* in the human brain. A naturally occurring form of gene overdose for both *APP* and *BACE2* is DS, caused by the trisomy of human chromosome 21 (T21) that harbours both *APP* and *BACE2* genes. As increased levels of soluble Aβ were observed already in foetal brains in DS^17^, we examined cerebral organoids grown from induced pluripotent stem cell (iPSC) generated by non-integrational reprogramming of primary cells donated by people with DS, including an isogenic DS (T21) iPSC model^18^, as a platform to analyse the T21-specific effects on APP proteolytic processing.

## Results

### Trisomy 21 (but not Dup*APP*) skews the ratios of Aβ non-amyloidogenic peptides

We compared organoids from isogenic iPSC lines, derived from the same individual with DS, mosaic for T21 and normal disomy 21 (D21) cells^18^. Cerebral organoids were derived following a standard protocol^19^, and shown to contain neurons expressing markers of all 6 layers of the human cortex (Supplementary Fig. 1) and no significant difference in the proportions of neurons and astrocytes between the D21 and T21 organoids (Supplementary Fig. 2). The integrity and copy number of the iPSC lines were validated at the point of starting the organoid differentiation, for chromosome 21 (Supplementary Fig. 3), and the whole genome (available on request).

T21/D21 status was further verified by interphase FISH on mature organoid slices, (Supplementary Fig. 4a). The C-terminal region of APP can be processed by the sequential action of different proteases to produce a range of protein fragments and peptide species, including Aβ (Supplementary Fig. 5). Aβ peptide profiles were analysed from organoid-conditioned media (CM) whereby each CM sample was taken from a 6cm dish culturing a pool of 12-16 organoids derived from one iPSC line, in total: n=15 CM samples for Exp1 (3 trisomic isogenic lines, 2 disomic isogenic lines, 3 timepoints each), n=12 CM samples for Exp2 (2 trisomic isogenic lines, 2 disomic isogenic lines, 3 timepoints each) and n=20 CM samples for Exp3 (1 trisomic isogenic line, 1 disomic isogenic line, 1 DupAPP line, 1 line each for two different unrelated DS individuals, 4 timepoints each). CM was collected at a timepoints between days 100-137 of culturing and analysed using immunoprecipitation in combination with mass spectrometry (IP-MS)^20^. Please see “Methods” and “Supplementary Data” sections for more detailed explanations, and statistical controls used for individual iPSC line-to-line comparisons. (Fig. 1a). Relative ratios were calculated of areas under the peak between the peptides of interest within a single mass spectrum (raw data example in Supplementary Fig. 6d), therefore unaffected by the variability in the total cell mass between wells growing organoids. The proportions of non-amyloidogenic peptides with the signature of BACE2 cleavage products, both as a putative θ-secretase (as reflected by the Aβ1-19 product) and putative AβDP or Aβ-clearance products (Aβ1-20 & 1-34), or combined, (relative to the sum of Aβ amyloidogenic peptides (Aβ1-38&1-39&1-40&1-42)) were approximately doubled in CM from T21 organoids, compared to isogenic normal controls, and reached levels of >80% of the amyloidogenic peptide levels (Fig. 1a). This result was fully reproduced in 3 independent experiments, each starting from undifferentiated iPSCs (3 vertical columns of graphs in Fig. 1a). In experiment 3, more recently generated iPSC lines from different individuals were introduced; from a euploid patient with FEOAD caused by Dup*APP*^21^, and from 2 unrelated people with DS (Supplementary Figs. 1-3). The 1-34&1-20/amyloidogenic ratios were not significantly different between D21 and Dup*APP* lines, suggesting the third copy of the *APP* gene alone did not cause any change in this ratio. Ratios of 1-34&1-20/amyloidogenic peptides and combined BACE2-products/amyloidogenics were significantly increased in T21 lines (combining all 3 T21 individuals) compared to D21 or Dup*APP* lines (Fig. 1a). The ratio of 1-19/amyloidogenics was significantly higher in T21 lines from the isogenic model, compared to its disomic isogenic control, and compared to Dup*APP*, but it was unchanged in the other two unrelated DS iPSC lines (see also Supplementary Information for a more detailed explanation). As the proportions of BACE2-unrelated α-site cleavage products (1-16, 1-17) were not different between T21 and isogenic D21 organoids (in any of the 3 experiments) (Fig. 1a), it can be predicted that the increased presence of 1-19, 1-20 and 1-34 peptides in T21 contributes towards an overall increase in soluble peptides that are non-amyloidogenic. The validity of this prediction was tested by an independent biochemical method (ELISA), by measuring the Aβ-peptide concentrations within the isogenic T21:D21 organoid CM comparison, which showed an increase in absolute concentrations caused by T21 for each Aβ 1-38, 1-40 and 1-42, with no difference in the Aβ 1-42/1-40 ratio between T21 and isogenic D21 lines, mirroring the readout in the absolute levels of IP-MS peaks (Supplementary Fig. 6). Analysis of IP-MS area under peak (used in Fig. 1 to calculate relative ratios) showed a near linear correlation when plotted against absolute peptide concentrations measured by ELISA, for each Aβ 1-38, 1-40 and 1-42 (Supplementary Fig. 6), validating our relative ratio calculations by an independent biochemical method.

**Fig. 1.**
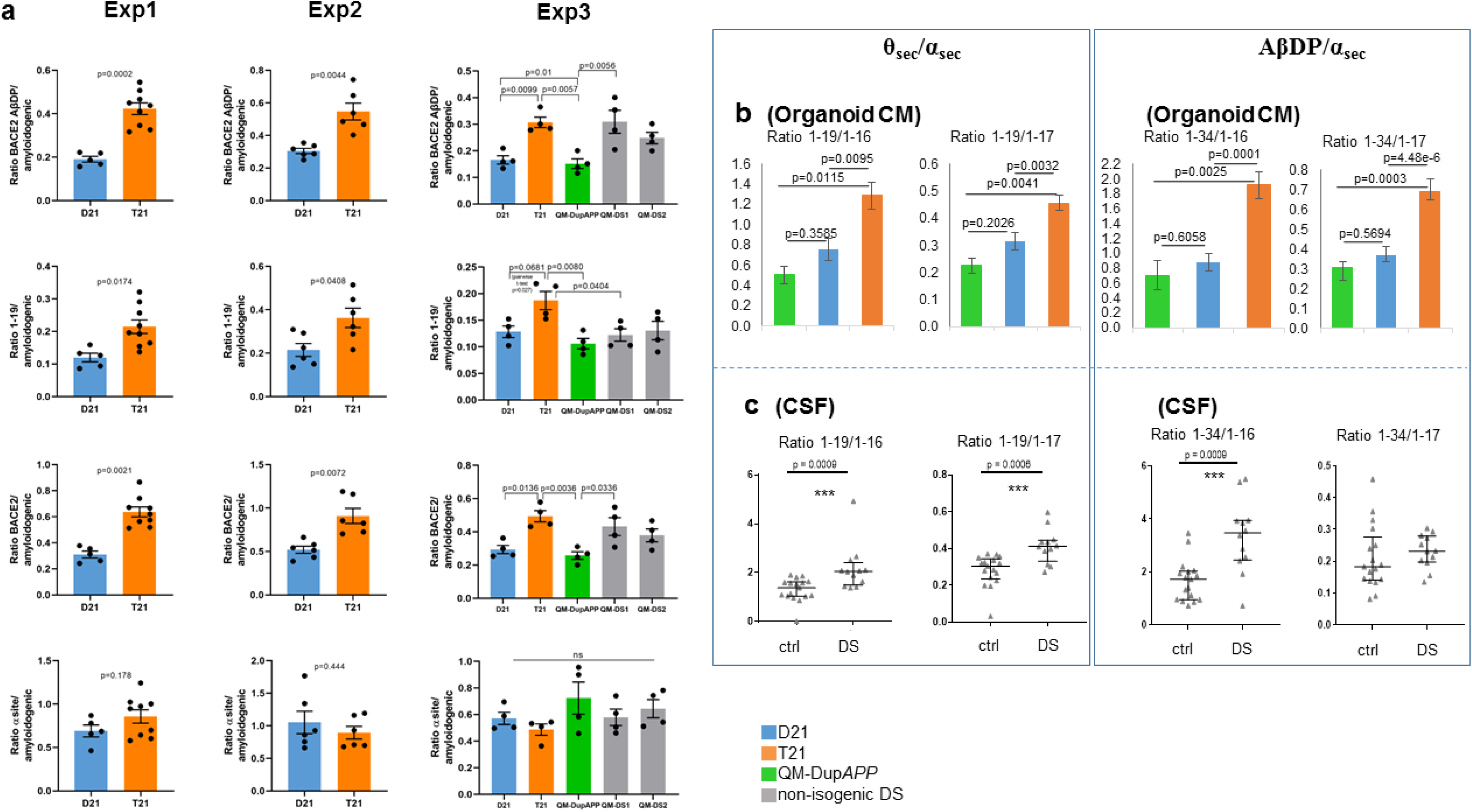
Aβ-peptide profiles secreted by trisomy-21 cerebral organoids. **a** Using Aβ IP-MS spectra from organoid (see Extended Data Figs. 1, 2, 4) conditioned media (CM), ratios were calculated of areas under the peak between the non-amyloidogenic and amyloidogenic peptides within a single mass-spectrogram. IP-MS spectra were produced for 3 timepoints (4 timepoints in exp3) for each iPSC-derived organoid line, in each of 3 independent experiments (each experiment starting at the point of undifferentiated iPSC). The team performing the IP-MS analysis was blinded to the genotypes in all experiments. BACE2-AβDP (clearance) products=[1-20&1-34], total BACE2=[1-19&1-20&1-34], amyloidogenic peptides=[1-38&1-39&1-40&1-42], α-site products=[1-16&1-17]. Exp1 and Exp2 p-values: Holm-Bonferroni sequential corrections (α=0.05) of 2-tailed student t-test comparisons. Exp3: Holm-corrected p-values after one way ANOVA. Error bars: standard error. Combined data for the isogenic iPSC lines for all 3 experiments passed the Holm-Bonferroni correction (α=0.05) of sequential 2-tailed student t-test comparisons of each peptide ratio shown in Fig1a (available on request). T21 and D21: isogenic iPCS derived from a single mosaic individual with DS published previously (Murray A et al. 2015), QM-DS1 and QM-DS2: unrelated DS iPSC, DupAPP: FEOAD iPSC. **b** All 3 experiments in Fig. 1a were combined to calculate the ratios of BACE2-related non-amyloidogenic peptides (1-19 or 1-34) to BACE2-unrelated non-amyloidogenic peptides (1-16 or 1-17) in organoid CM. Holm-corrected p-values after one way ANOVA are shown. Error bars: standard error. **c** Same ratios as in part A were calculated on IP-MS spectra obtained from cerebrospinal fluid samples of people with DS and age-matched normal controls. Data are presented as mean ± 1SD.

To estimate the contribution of BACE2 towards the anti-amyloidogenic pathway relative to other anti-amyloidogenic cleavages at the α-site, we calculated the peptide ratios of 1-19/1-16 or 1-17 (θ secretase/α secretase products) and 1-34/1-16 or 1-17 (BACE2 AβDP/ α secretase products). We observed that T21 organoids produce statistically highly significant increases in all four of these ratios, relative to isogenic D21, or non-isogenic Dup*APP* organoids (Fig. 1b). Therefore, we conclude that T21 causes these effects in our organoid system. The D21 ratios were not significantly different to Dup*APP*, suggesting that the third copy of genes other than *APP* causes these effects, though this needs to be tested on a larger number of individuals.

As the peptide profiling data strongly favour the hypothesis of a genetic-dose-sensitive anti-amyloidogenic action of BACE2, we sought to zoom in on the *BACE2* genetic locus in a systematic SNP-array analysis of 554 individuals recruited through the LonDownS Consortium who had undergone detailed assessment for dementia^22, 23^; 93 single nucleotide polymorphisms (SNPs) located within the *BACE2* locus +/-50kb, were genotyped, and dementia age-of-onset determined, as described in Methods. We detect two new BACE2 SNPs (purple, Supplementary Fig. 7) correlating with age of dementia onset in the DS cohort of the LonDownS Consortium, located in close proximity to a previously reported SNP (red, Supplementary Fig. 7) ^24^. All of these 3 SNPs cluster in <4kbp segment, which is fully contained within a 12kbp deletion (blue line, Supplementary Fig. 7) that caused a *de novo* EOAD in a euploid patient^25^ (Supplementary Fig. 7). These data corroborate the notion that subtle genotypic variation in *BACE2* levels may play an important role in affecting the age of dementia onset in both DS and non-DS individuals.

### Non-amyloidogenic Aβ peptide ratios mirror between T21 organoids and DS-CSF

In order to assess if the peptide ratio differences from Fig. 1b have any relevance *in vivo*, we analysed the Aβ-peptide profiles immunoprecipitated from human cerebrospinal fluid (CSF). We have previously produced IP-MS data on CSF from people with DS and age-matched controls^26^. We repeated the calculations shown for organoids in Fig. 1b, on IP-MS results from CSF samples from DS (n=17) and age-matched euploid people (n=12). All four relative ratio calculations showed an increase in peptide ratios in CSF from people with DS, compared to age-matched euploid controls, of which three comparisons were statistically highly significant (Fig. 1c). This suggests that in DS brains, the third copy of *BACE2* skews the anti-amyloidogenic processing significantly towards BACE2-cleavages, relative to other anti-amyloidogenic enzymes cleaving at the α-site. Importantly, these CSF results validate the *in vivo* relevance of the peptide ratios obtained using CM from iPSC-derived cerebral organoids (comparison of Fig. 1b and Fig. 1c).

### Aβ-degrading activity of BACE2 is cross-inhibited by clinically trialed BACE1-inhibitors

Chemical inhibition of BACE1 remains an attractive therapeutic strategy for AD. As BACE2 is a homologous protein, most inhibitors tested in clinical trials also cross-inhibit the (pro-amyloidogenic) β-secretase activity of BACE2, which has been proven as the cause of several unwanted side-effects, such as skin pigmentation changes. As our data suggest that the opposite, Aβ-degrading, activity of BACE2 plays an important role, we designed a new FRET-based *in vitro* assay, in which efficient AβDP-cutting after Aβ aa34 by BACE2 at pH=3.5 could be measured (Fig. 2), while zero activity by BACE1 was detectable under same conditions (Supplementary Information). We demonstrated that at least two BACE1 inhibitor compounds (of which one recently used in clinical trials) inhibit the AβDP (Aβ-clearance) activity of BACE2 in a dose-dependent manner (Fig. 2). This has, to our knowledge, so far not been shown, and could provide an additional explanation for the failure of some BACE1-inhibitor clinical trials, and should be taken in consideration when testing new inhibitors.

**Fig. 2.**
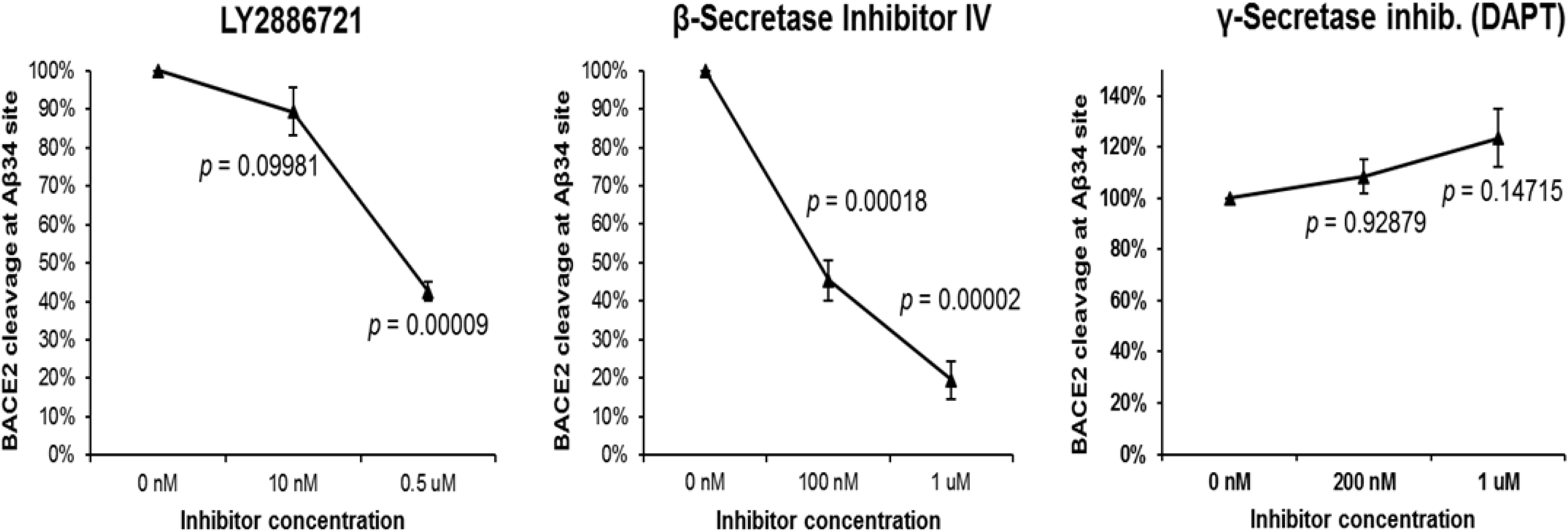
FRET based assay for BACE2 cleavage. A newly custom designed FRET reagent (spanning the Aβ34 site) was digested at pH=3.5 by the human BACE2 in presence or absence of the stated inhibitors for 2h. Enzyme activity was defined by measuring the fluorescence increase before and after the incubation. Blank-subtracted fluorescence units were normalized to the control digest and a one-way ANOVA was performed. P-values were calculated with a post-hoc Bonferroni multiple comparison (only pairs relative to the untreated control simultaneously compared). Error bars: standard error.

### AβDP product (Aβx-34) colocalises with BACE2 in human brain and organoid neurons

As *in vitro* experiments showed that BACE2 can very efficiently cleave the Aβ34-site in the FRET peptide (Fig. 2) and synthetic Aβ1-40 peptide in solution at an acidic pH^12^, we sought to visualize if the presence of the substrate (Aβ1-40), enzyme (BACE2) and one of the products of this reaction (Aβ1-34) can be detected in our organoids, in a sub-cellular compartment known to be acidic. Firstly, by immunofluorescence (I.F.) using pan-anti-Aβ (4G8), anti-BACE2, or neo-epitope-specific antibodies against Aβx-40 and Aβx-34^27^, we detected significantly higher signals (normalized to pan-neuronal marker) in T21 organoid neurons, compared to isogenic D21 ones (Supplementary Fig. 4b-d). Pearson’s coefficient showed a high level of colocalisation (>0.55) of both the main substrate (Aβx-40) and its putative degradation product (Aβx-34) with BACE2 in neurons of cerebral organoids, in LAMP2+ compartment (known to be a subset of lyzosomes, therefore low pH vesicles) (Fig. 3 & Supplementary Fig. 8). In comparison, the Pearson’s coefficient for BACE1 with Aβx-34 was only 0.16 (Fig. 3 & Supplementary Fig. 8), and its pattern of sub-cellular localization was different to BACE2 (high colocalization with Rab7 and Sortilin, much lower with LAMP2). Using I.F. on human brain sections, a similar highly significant difference was observed (Fig. 4a, b): Aβx-34 colocalised with BACE2 (0.52 (±0.034SEM)) as opposed to BACE1 (0.01 (±0.021SEM)). The colocalised signal of Aβx-34 and BACE2 was seen in 3 categories of objects (Fig. 4), in all analysed samples: 4 individual DS-AD brains (Fig. 4a-c), 5 euploid sporadic AD subjects (example in Supplementary Fig. 14a, for complete list of brain samples see Supplementary Table 1) and (in the fine vesicle compartment only) in 5 non-demented control euploid subjects’ neurons (age 42-84), as well as DS brain from a 28yr old with no plaques or dementia, (examples in Fig. 4d, for complete list of brain samples see Supplementary Table 1). Lambda scanning and Sudan black B stainings were independently used to subtract the autofluorescence of lipofuscin granules (Supplementary Fig. 15f, g). This has proven that the fine vesicular pattern and large amorphous extra-cellular aggregates are not autofluorescent lipofuscin granules, but real colocalisations of BACE2 and Aβx-34 (Supplementary Fig. 15). Colocalised signals of Aβx-34 and BACE2 were particularly strong in areas surrounding neuritic plaques (Fig. 4a-c).

**Fig. 3.**
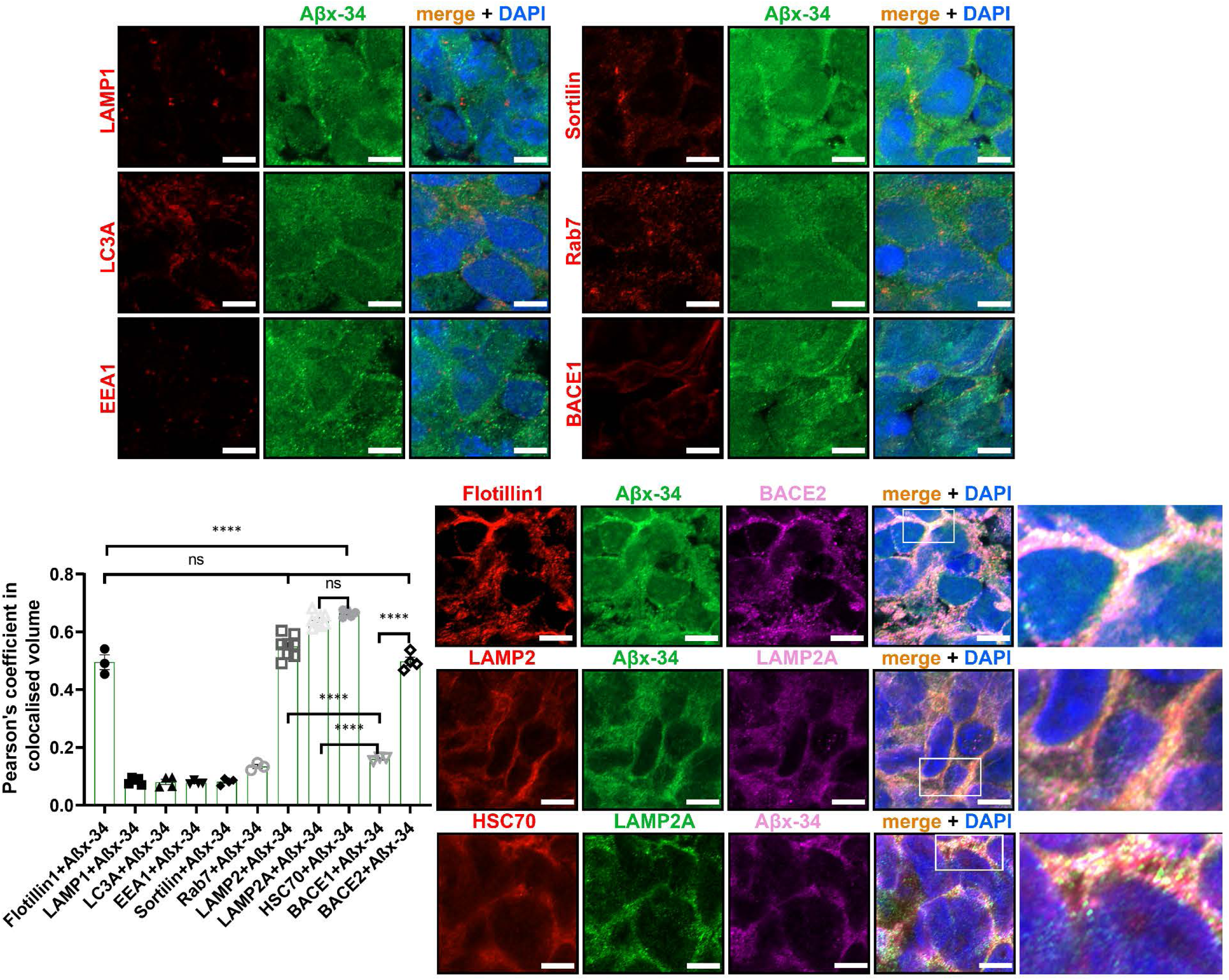
Sub-cellular compartment localisation of Aβ degradation product Aβx-34 in hiPSC-derived cerebral organoid sections. Pairwise Pearson’s coefficient of colocalisation for a pair of co-stained antibodies: Aβx-34, and a specific marker for the sub-cellular vesicle compartment: lipid rafts (Flotillin1), lysosomes (LAMP1), macro-autophagosomes (LC3A), early endosomes (EEA1), macro-autophagosome-lysosome fusion/exosomes (Sortilin), late endosomes (Rab7), specific sub-sets of lysosomes (LAMP2, LAMP2A) and CMA-chaperone (HSC70). In the final two columns of the histogram, the Pearson’s colocalisation level was shown between Aβx-34 and BACE1 or BACE2, respectively (repeated in more detail in Supplementary Figure 8). Representative images of the organoid stainings from which the coefficients were calculated are shown in the panels. Last column in the bottom right panel is the zoomed-in inset from the previous column. Images were captured using Airy-Scan Zeiss confocal microscopy, and single 0.16μm slices are shown (from 20μm full z-stack analysed). Error bars: standard error, p-values: after standard one-way ANOVA using post-hoc Bonferroni multiple comparison calculation. Scale bar: 5μm.

**Fig. 4.**
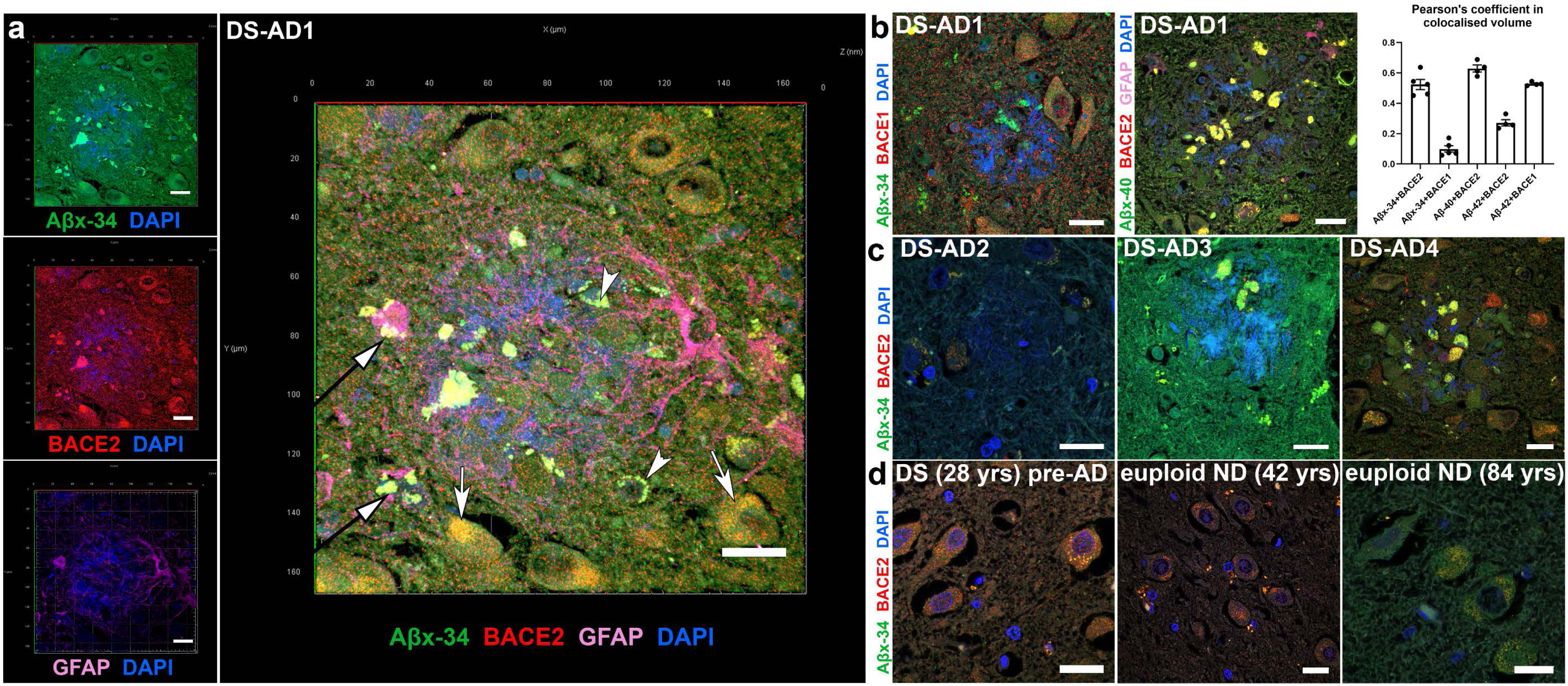
Localisation of AβDP degradation products and Aβ peptides with BACE2 in hippocampal sections of the human post-mortem brain. **a** Immunofluorescence analysis of the brain of DS-AD-1 co-stained for Aβx-34, BACE2 and GFAP. A typical near-circular neuritic plaque is shown (in which DAPI faintly stains the fibrillar amyloid deposits). Arrows indicate 3 categories of objects in which the colocalisation of BACE2 and the AβDP product Aβx-34 is observed. White arrows: intra-neuronal fine-vesicular pattern; White arrowheads: large intra-neuronal spherical granules (lipofuscin); Black arrows with white arrowheads: amorphous extra-cellular aggregates. See Methods and FigS6 for experiments controlling the extent of lipofuscin autofluorescence effects. **b** DS-AD1 brain co-stained for Aβx-34 and BACE1, or Aβx-40, BACE2 and GFAP, and Pearson’s coefficient of colocalisation for proteins stained in parts A and B, with the addition of the staining for Aβx-42 neo-epitope (not shown). Error bars: SEM. **c** Same I.F. staining combinations as in part A (except for GFAP) were used in 3 additional brain samples: DS-AD-2, 3 and 4. **d** brain sample co-stained for BACE2 and Aβx-34 of a 28yrs old person with Down syndrome without dementia, and euploid non-demented (ND) controls aged 42 and 84. Scale bar: 20μm.

**Fig. 5.**
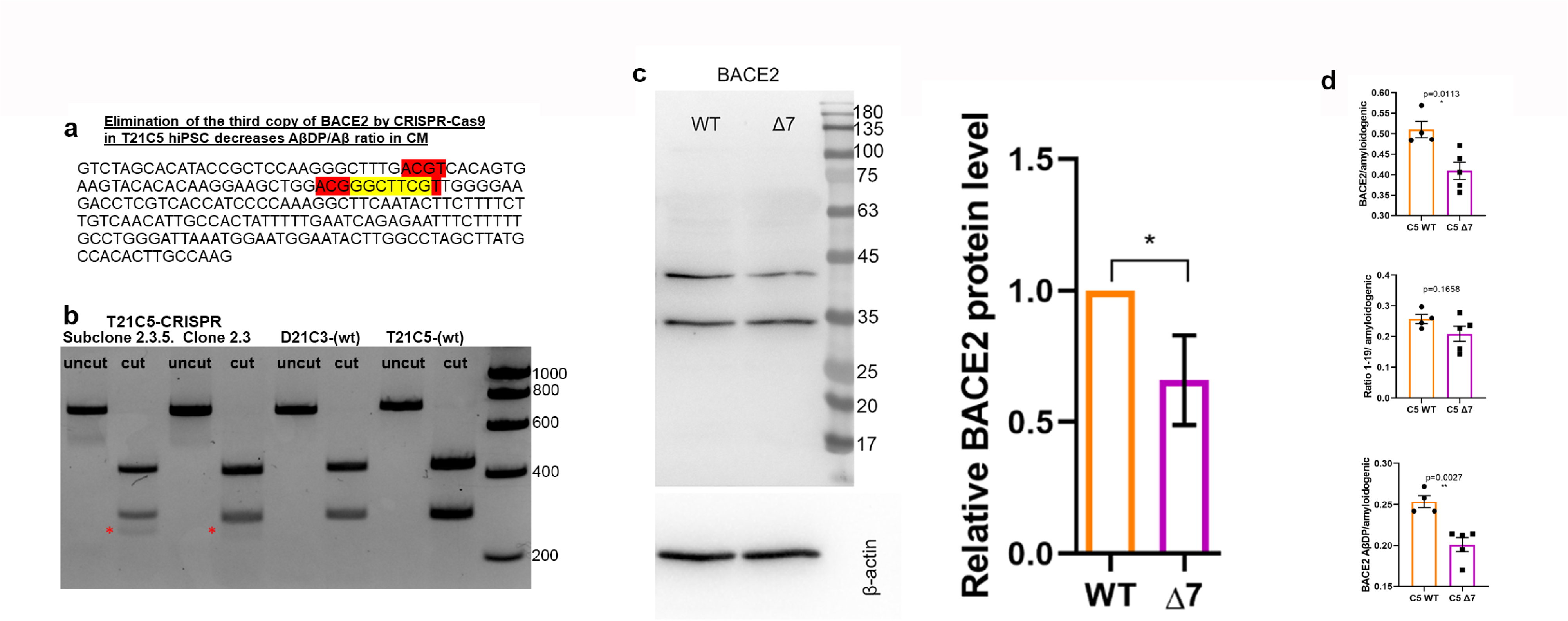
CRISPR/SpCas9-HF1-mediated reduction of BACE2 copy number from 3 to 2 in the T21C5 hiPSC line. **a** *BACE2* exon3 sequence with 7bp deletion (yellow) provoked by the CRISPR/SpCas9-HF1 is shown. Red: restriction endonuclease HpyCH4IV sites (a de novo HpyCH4IV site is generated by the 7bp deletion). **b** agarose gel electrophoresis of the 733 bp PCR product containing the targeted site before (uncut) and after digestion with HpyCH4IV(cut), for the initial clone 2.5, and its colony-purified sub-clone 2.3.5 (renamed further below as “Δ7”). The 294bp fragment in 2.3.5 is reduced to 65% of the wt value (normalized to the 439bp band), and a *de novo* 255 bp fragment appears in CRSPR targeted line (red asterisk). **c** Western blot stained with anti-BACE2 antibody of the lysates of the iPSC line Δ7 compared to the wt T21C5 iPSC line. Quantification of the total actin-normalised BACE2 signal showed a significant reduction in Δ7 compared to T21 unedited line. Error bars: standard error, p-value: student’s t-test. **d** BACE2 AβDP/amyloidogenic peptides ratio after IP-MS analysis of CM produced by the 48DIV organoids derived from the iPSC line Δ7 compared to the T21C5wt control were significantly decreased. Error bars: standard error, p-values: two-tailed t-test comparison.

As AβDP cleavage by BACE2 is efficient only at low pH, we sought to analyse in more detail the BACE2 and Aβx-34 co-localisation in highly acidic cellular compartments. For this reason, we costained lysosome markers LAMP1 or LAMP2 with Aβx-34. Additionally, macro-autophagic vacuoles containing Aβ were shown to accumulate in AD distended neurites^28^, which is why we also stained with the macro-autophagosome marker LC3A. As we further found that Aβx-34 did not colocalise with LAMP1 or LC3A, but colocalised strongly with LAMP2 (Fig. 3, Supplementary Figure 8 and Supplementary Information), we tested colocalisation with the components of an alternative autophagy pathway: chaperone-mediated autophagy (CMA), and found a very high level of colocalisation (Fig. 3).

### Trisomy of *BACE2* skews non-amyloidogenic Aβ peptide ratios and suppresses AD-like pathology in organoids

Using CRISPR/SpCas9-HF1, we eliminated a single copy of *BACE2* in the trisomic iPSC line C5 (T21C5Δ7, a Δ7bp in *BACE2* exon3, knocking out 1 of 3 copies of the gene), while maintaining the trisomy of the rest of chromosome 21 (Fig. 5a-c, Supplementary Fig. 9, Supplementary Information). Total actin-normalised BACE2 signal showed a 27%-34% reduction in Δ7 compared to T21 unedited line, and no significant difference compared to D21 control (Fig. 5c, Supplementary Fig. 10). Total protein level of APP in Δ7 remained at trisomic levels, significantly increased compared to the disomic control (Supplementary Fig. 10). The CRISPR correction of *BACE2* gene dose from 3 to 2, resulted in a significant decrease in levels of putative BACE2-AβDP (Aβ-clearance) products (1-20&1-34), as well as total BACE2-related non-amyloidogenic peptides (1-19&1-20&1-34), relative to amyloidogenic peptides (Fig. 5d).

This pinpoints the triplication of *BACE2* as a likely cause of specific anti-amyloidogenic T21 effects we observed in Fig. 1a. Furthermore, we used two different dyes to detect any presence of amyloid deposits (the traditional Thioflavine S, and a newer, more sensitive dye AmyloGlo^29^) in organoid sections. Remarkably, elimination of the third *BACE2* copy caused the T21 organoids (that had not shown any overt amyloid deposits at 100DIV, see T21C5 in Supplementary Fig. 11, top row) to develop extremely early AD-plaque like deposits (AmyloGlo+ and Thioflavine S+) in the cortical part of the organoid by 48DIV (Supplementary Fig. 11, middle row), that progressed aggressively and became much stronger and denser by 96 DIV, accompanied by massive cell death (Supplementary Fig. 11, bottom row, Supplementary Fig. 12).

In order to prove that extracellular deposits staining positively with amyloid dyes really are related to hyperproduction of Aβ amyloidogenic peptides, we cultured T21C5Δ7 organoids in media containing high concentrations of β and γ secretase inhibitors. Early T21C5 and T21C5Δ7 organoids were treated with a combination of β-secretase inhibitor IV and compound E (γ secretase inhibitor XII) (Supplementary Table 2) from 20DIV to 41DIV (Fig. 6). Amyloid-like deposits were readily detected with AmyloGlo in the untreated and vehicle only treated T21C5Δ7 organoids (Fig. 6b), but were completely absent from T21C5Δ7 organoids treated with β and γ secretase inhibitors. Inhibitor treatment also significantly reduced the number of neurons expressing pathologically conformed Tau (TG3-positive cells) in the T21C5Δ7 compared to untreated controls (Fig. 6c). No AmyloGlo positive aggregates or TG3-positive cells were detected in T21C5 organoids under any treatment conditions at DIV41 (Fig. 6a, c) and were also absent in the same organoids at DIV100 (Fig. 7 g, l, Supplementary Fig. 11). Also, no obvious deleterious effects of the inhibitors, or vehicle control, could be seen in early unedited T21C5 organoids.

**Fig. 6.**
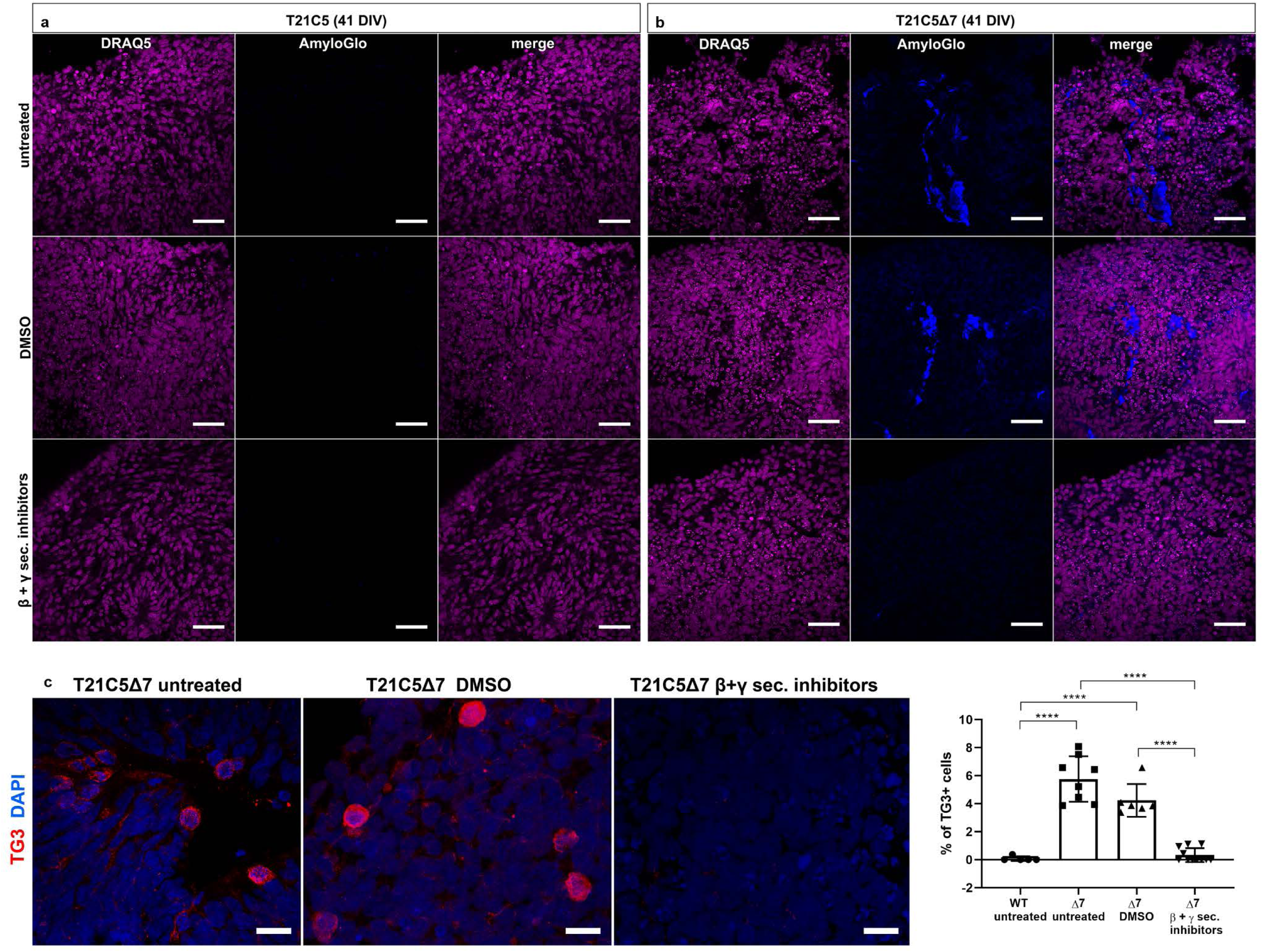
CRISPR/SpCas9-HF1-mediated reduction of BACE2 copy number from 3 to 2 in the T21C5 hiPSC line provoked early AD-like pathology in organoids. a-c. Early AD-like pathology was provoked in 41DIV T21C5Δ7 organoids, but was not detected in T21C5 parental organoids. -**a-b** Treatment of the T21C5Δ7 with combined βI-IV (β-sectretase inhibitor) and Compound E (γ-secretase inhibitor) from 20-41DIV completely prevented the formation of extracellular amyloid deposits. Staining with amyloid specific dye (AmyloGlo) and nuclear dye (DRAQ5). Scale bar 50μm. **c** β- and γ-secretase inhibitor treatment highly significantly reduced the presence of TG3+ (pathologically conformed Tau) cells in T21C5Δ7 organoids compared to untreated T21C5Δ7 organoids. Scale bar 20μm. Error bars: SD, ****p<0.0001. Only statistically significant differences are shown.

**Fig. 7.**
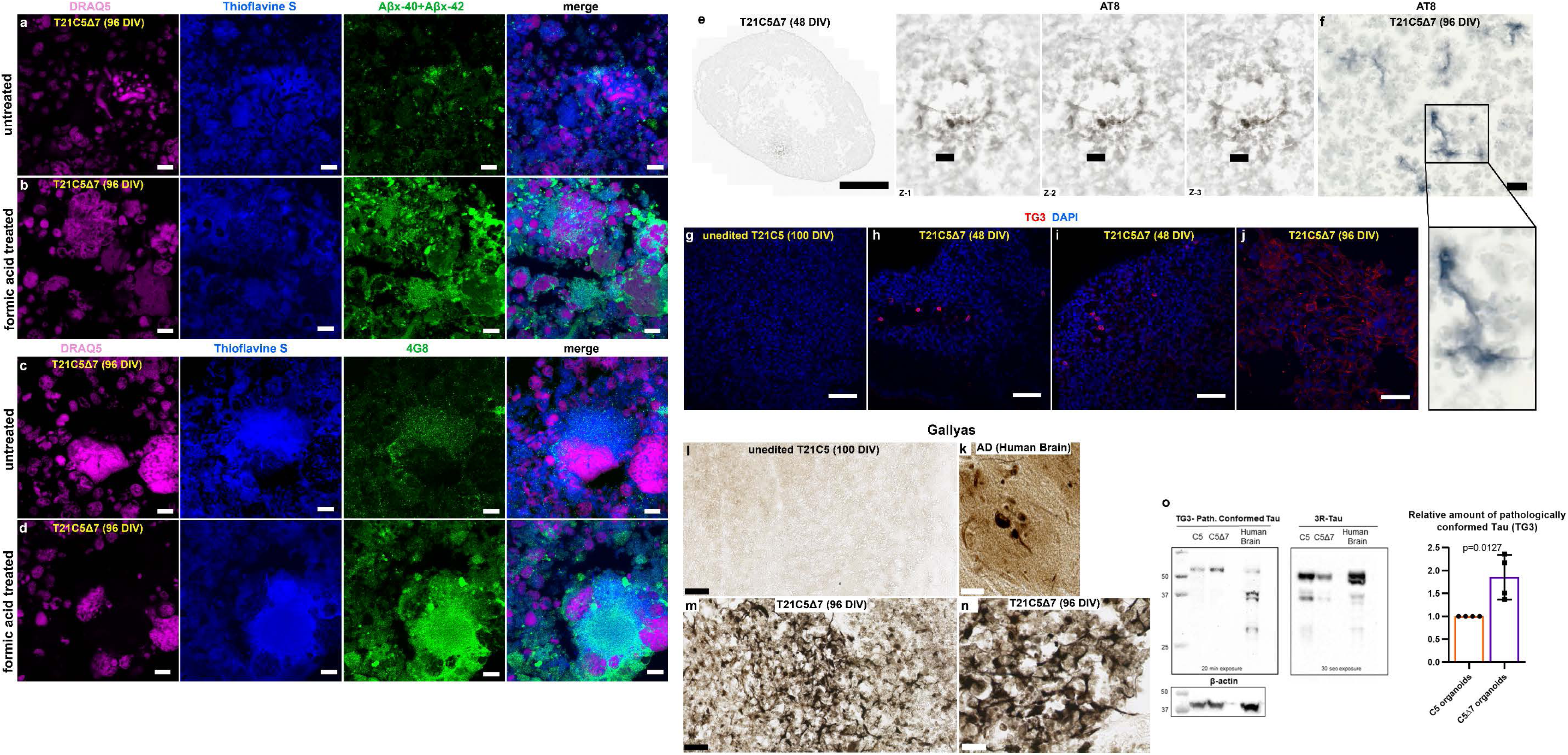
Amyloid and Tau pathology are shown with six different methods in T21C5Δ7 organoids that have BACE2 copy number reduced from 3 to 2 by CRISPR/Cas9. a-d. The signal of amyloid specific antibodies Aβx-40 + Aβx-42 (a, b), or 4G8 (c, d) colocalising with Thioflavine S in T21C5Δ7 (96DIV) organoids was drastically increased upon treatment with 87% Formic acid for 10 minutes at RT, proving it contains the insoluble extracellular β-amyloid deposits. Scale bar: 10μm. **e** AT8 (hyper-phosphorylated Tau) positive neurites within plaque-like structure in 48DIV organoids. Left: the whole organoid slice, scale bar: 500μm. Right: zoom in on the plaque-like structure from E, in the three individual z-slices (interval between slices, 1μm; scale bar: 20μm). **f** AT8 (hyper-phosphorylated Tau) positive neurites in 96DIV organoids. Scale bar: 10μm. **g-j** TG3 (conformationally altered Tau) staining of unedited control T21C5 (100DIV) (g), CRISPR-edited T21C5Δ7 (48DIV) with TG3 positive neurons in 48DIV organoids (h, i), and CRISPR-edited T21C5Δ7 (96DIV) showing many TG3+ neurons with diffuse staining of extracellularized mal-conformed Tau aggregates (j). Scale bar: 50μm. k-n Gallyas staining of human AD brain (k), unedited control T21C5 (100DIV) (l), CRISPR-edited T21C5Δ7 (96DIV) (M,N) shows negative staining in parental unedited organoid (l) and very strong signal in neurons and plaque-like associated neurites within T21C5Δ7 organoid (m, n). Scale bars: 50μm (l, m) and 20μm (n) and 5μm (k). **o** Representative Western blot of T21C5 and T21C5Δ7 organoid lysates stained using antibodies against pathologically conformationally altered Tau (TG3) or general 3 repeat (3R) Tau. β-actin was used as a loading control. Human brain tissue of a 75 year old is shown for comparison. Comparison of the average values (n=4) for CRISPR-edited T21C5Δ7 showed a highly significant relative increase in TG3 compared to unedited (n=4) T21C5 organoids, as indicated in the graph, p=0.0127.

Further histo-pathological verification showed that elimination of one copy of *BACE2* triggered progressive accumulation of extracellular deposits that co-stain with Thioflavine S and antibodies against Aβ, both 4G8 and neo-epitope specific Aβx-40&Aβx-42. The antibody signal intensity in colocalisations with Thioflavine S drastically increased upon pre-treatment with 87% formic acid (Fig. 7a-d), proving that the deposits contain insoluble Aβ material. This is further corroborated by the isolation of fibrillary material from the detergent-insoluble fraction of the CRISPR-edited organoid. When viewed by Transmission Electron Microscopy (TEM) the filaments found exhibited a straight morphology of <10nm diameter (Supplementary Fig. 13a), closely resembling fibrils grown *in vitro* from synthetic Aβ1-40 peptide (Supplementary Fig. 13c). Furthermore, neuritic plaque-like features were detected by IHC co-staining with Gallyas in CRISPR-edited organoids (Fig. 7m, n), but not their unedited T21 control (Fig. 7l). Human brain from an AD patient is shown for comparison stained with Gallyas (Fig. 7k). Tau pathology was also observed by IHC using the hyper-phosphorylated Tau antibody AT8 (Fig. 7e, f), and by I.F. for conformationally altered Tau (TG3, Fig. 7g-j). The relative increase in the amount of conformationally altered (pathological) Tau in CRISPR-edited organoids T21C5Δ7, compared to unedited T21 control organoids, was also independently confirmed by immunoblotting using TG3 antibody. As shown in Fig. 7o, the protein material isolated from T21C5Δ7 organoids produced significantly more TG3 signal than unedited controls, albeit having a weaker signal with the general 3R-Tau antibody (consistent with the observed neuronal loss, Supplementary Fig. 12).

### Alzheimer’s disease-like pathology develops reproducibly in unedited cerebral organoids from 71% of DS donors, and it is donor-specific

Our data in Figs. 5, 6, 7 show that severing the *BACE2* dose by a third, using CRISPR/Cas9, might tip the balance against the anti-amyloidogenic activity, and provoke AD-like pathology. Our data in Fig.1 suggest that anti-amyloidogenic activity of BACE2 is gene-dose dependent, and its level varies between individuals, with SNP allelic differences in *BACE2* gene correlating with age of dementia onset. We therefore hypothesized that organoids grown from some people with DS may develop AD-like pathology without any CRISPR-Cas9 intervention. We then tested this hypothesis using iPSC lines from 6 different individuals with DS, and one Dup*APP* patient (Table 1). We detected amyloid-like aggregates (both diffuse and compact in appearance) in 5/7 unedited iPSC-derived organoids from people with DS, and one with Dup*APP* (Fig. 8).

**Fig. 8.**
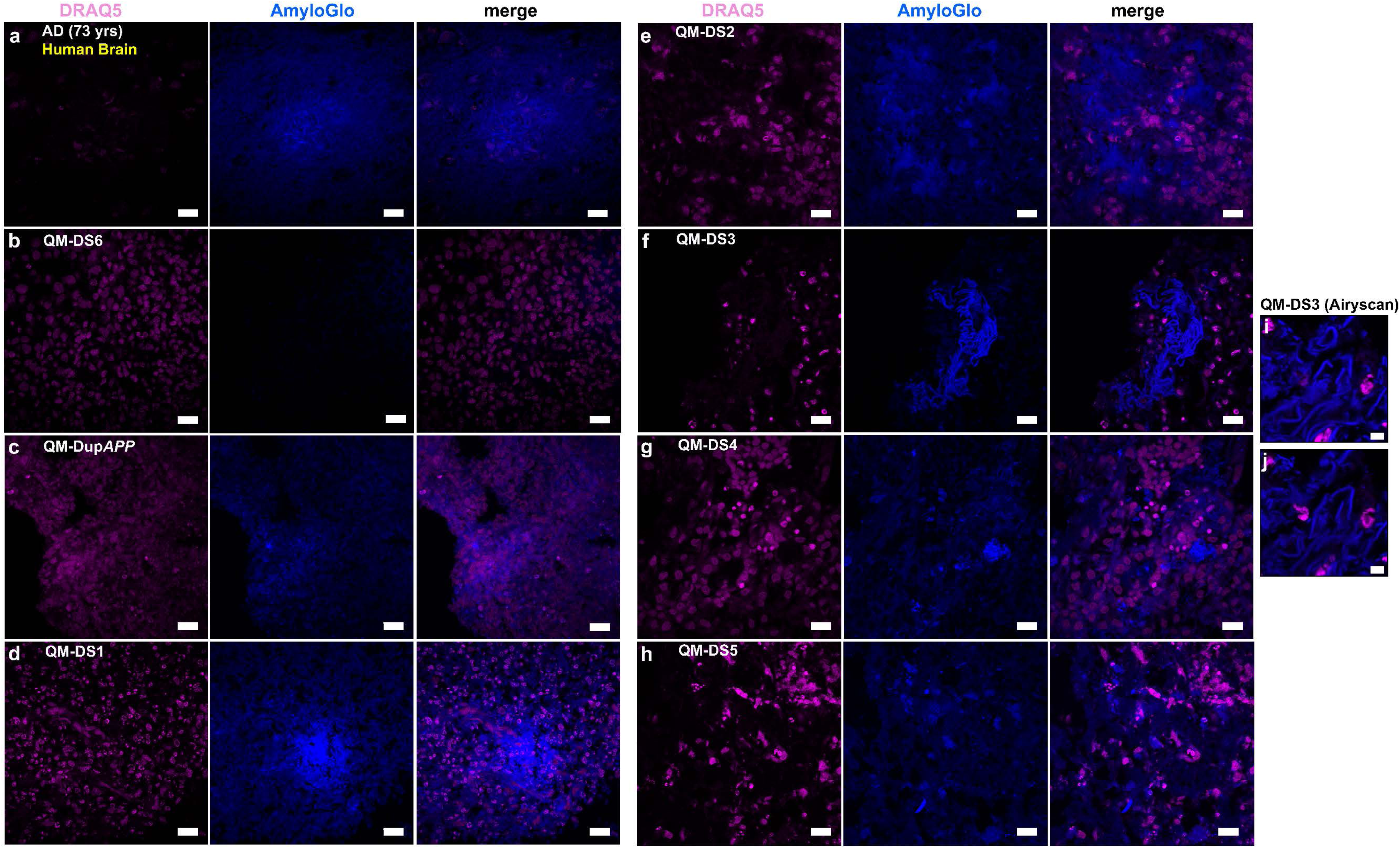
Amyloid-like pathology, staining with AmyloGlo, is shown in different lines of organoids and Human AD Brain served as positive control. **a** Human AD Brain (73 yrs) shows amyloid plaques in the Entorhinal cortex. **b** QM-DS6 (DIV100) shows no AD pathology. **c, d, e, f, g, h** QM-DupAPP and QM-DS1, 2, 3, 4, 5 show AmyloGlo positive aggregates, similar to Human Brain. Scale bar: 20μm. **i** and **j**: Airyscan analysis of QM-DS3 showing super-resolution images of AmyloGlo positive material with fibrillar-like appearance. Scale bar: 5μm.

**Table 1.**
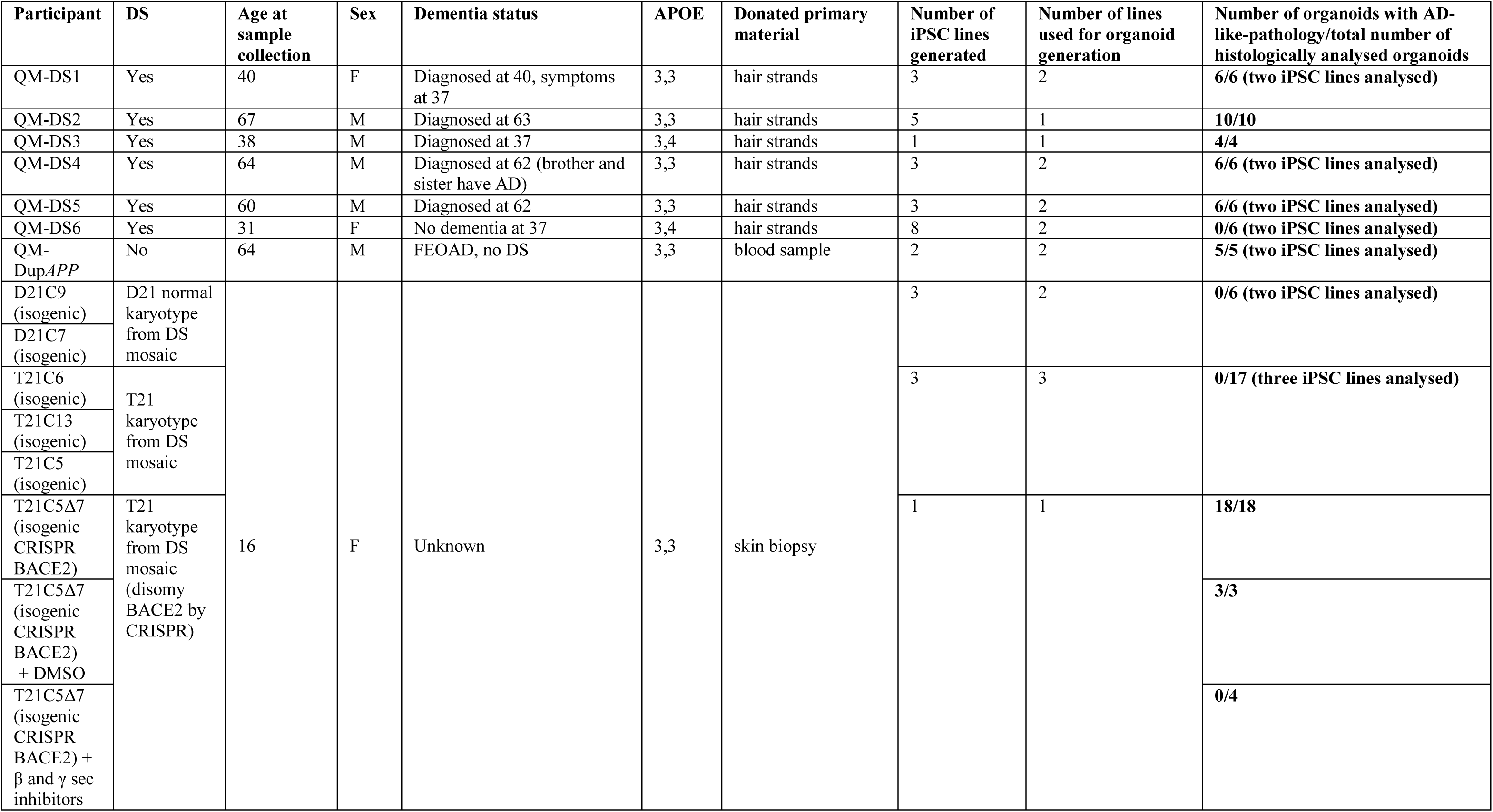
List of participant donors of cells for iPSC-organoid generation and reproducibility of AD-like pathology (by histological analysis)

The two donors whose iPSC-organoids did not show pathology are (i) the T21 iPSC from our isogenic model (whose clinical status is unknown) and (ii) QM-DS6, a donor who remains free from dementia symptoms at age 37 (Table 1). Organoids from another 5 DS donors, and one Dup*APP* patient, (all diagnosed with clinical dementia) all showed presence of diffuse and compact amyloid-like deposits (Fig. 8) as well as presence of neuritic plaque-like features (focal hyper-phosphorylated tau (AT8+), conformationally altered tau (TG3+) and filamentous Tau (AT100+)) within neuropil neurites within plaque-like circular foci (Fig. 9a-n). This was corroborated by Gallyas intra-neuronal positivity (Fig. 9o-t). Similarly as for T21C5Δ7, we were able to isolate fibrillary material from the detergent-insoluble fraction of QM-Dup*APP* organoid (Supplementary Fig. 13b), that on TEM resembled fibrils grown *in vitro* from synthetic Aβ1-40 peptide (Supplementary Fig. 13c). Most importantly: tested individual organoids from one donor (from multiple iPSC lines and multiple independent experiments) either all did (Dup*APP*, QM-DS1-5), or all did not (isogenic T21, QM-DS6) show AD-like pathology (Table 1), proving the pathology is donor dependent. This open possibilities of developing assays for pre-therapy risk-stratification and individualized drug-response quantitation.

**Fig. 9.**
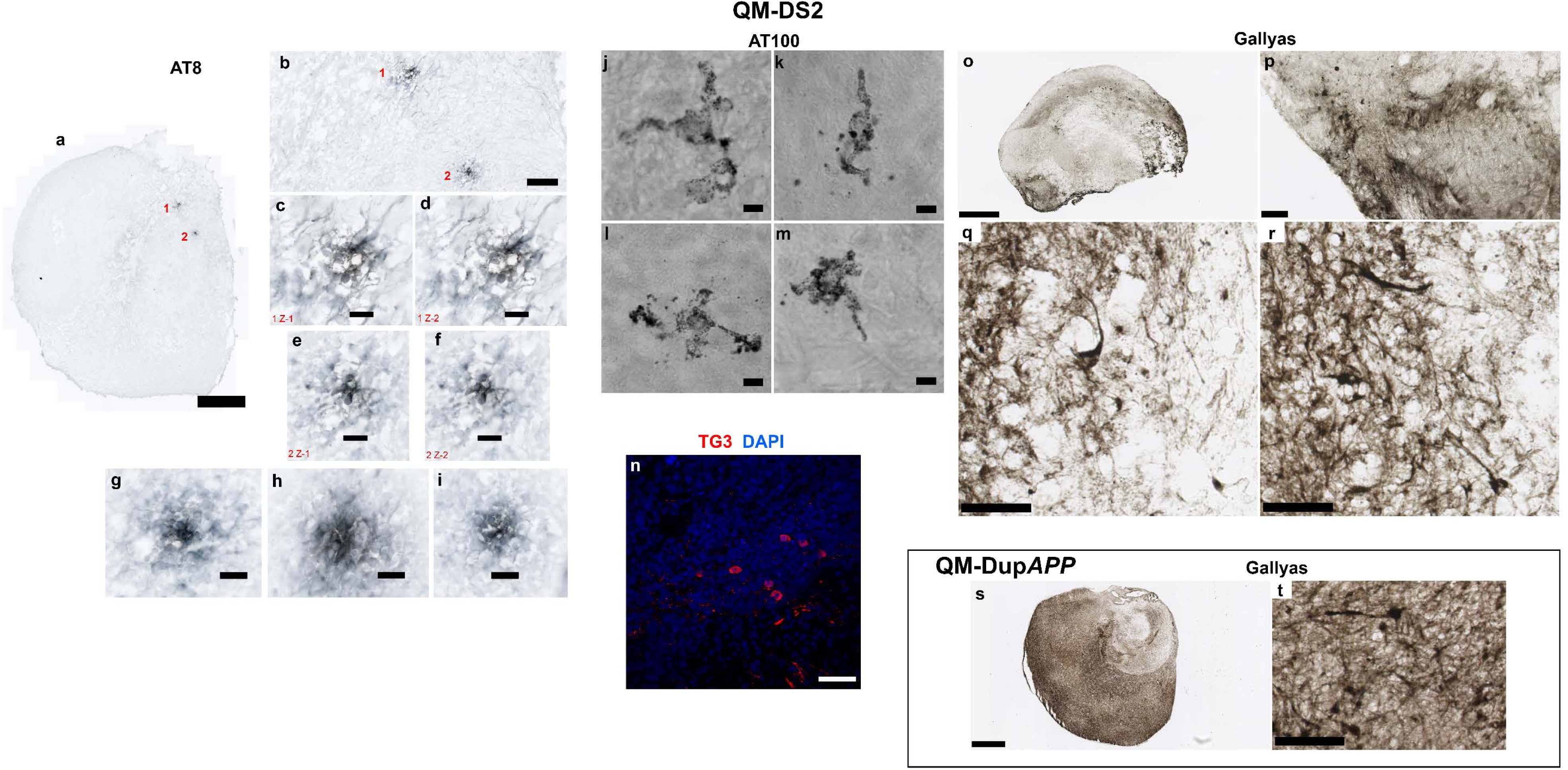
Tau pathology, staining with Gallyas, and with three different antibodies (hyper-phosphorylated, conformationally altered, and filamentous Tau) in QM-DupAPP (100DIV) and QM-DS2 (100DIV) organoids. **a** Scan of whole QM-DS2 organoid section shows two hyperphosphorylated Tau foci (neuritic plaque-like structures). Scale bar: 500μm. **b** Zoom in on the same foci. Scale bar: 100μm. **c** and **d** AT8 positive neurites in the pathological structure number 1 in two individual z-slices. Scale bar: 20μm. **e** and **f** AT8 positive neurites in the pathological structure number 2 in two individual z-slices. Scale bar: 20μm. **g-i** AT8 positive neurites in further three pathological foci found at a different depth, from the same organoid (not shown at lower magnification). Scale bar: 20μm. **j-m** AT100 (filamentous Tau) positive neurons in the cortical layer of QM-DS2 organoid, partly showing “ballooned neuron” pathology. Scale bar: 5μm. **n** TG3 (conformationally altered Tau) positive cells in the cortical layer of QM-DS2 organoid. Scale bar: 50μm. **o** Scan of whole QM-DS2 organoid section stained with Gallyas. Scale bar: 500μm**. p, q, r** zoom in on the parts of the same organoid shows strongly Gallyas-positive individual neurons. Scale bar: 50μm (**p**), 20μm (**q** and **r**). **s** Scan of whole QM-DupAPP organoid section stained with Gallyas. Scale bar: 500μm. **t** Zoom in on the same organoid shows equally strong individual neurons as in QM-DS2 organoid. Scale bar: 50μm.

## Discussion

Several human brain studies show detectable expression and β-secretase activity of BACE2, though at much lower levels than that of BACE1^30–33^. Chemical inhibition of β-secretase activity is an attractive therapeutic approach aimed at reducing the production of Aβ^34–36^. Complete knock-out of *BACE1* abolished all β-secretase activity in mouse neurons, while leaving some degree of β-secretase activity in astrocytes^37^. This activity was abolished by the complete knockout of both *BACE1* and *BACE2*, leading to a hypothesis that a BACE2-driven β-secretase activity in astrocytes may contribute to accelerate the Aβ-production and AD-pathology in DS^37^. In human brain, the β-secretase activity of BACE1 correlated positively with the amount of Aβ, whereas the β-secretase activity of BACE2 did not^30^. On the other hand, SNPs at the *BACE2* locus (and not *BACE1*) correlate with the age of onset of dementia in people with DS^24^, as well as sporadic LOAD in euploid people in the Finnish population^38^, and a recent report showed that a *de novo* intronic deletion within one allele of *BACE2* caused EOAD in a 50 year old euploid person^25^. All of the above data (and new data we show in Supplementary Fig. 7) implicate that a single allele alteration in the genetic dose of *BACE2* is capable of affecting the risk of AD-dementia, but do not resolve the question whether BACE2 *per se* acts predominantly as an accelerator, or a suppressor of AD pathology. The answer to this question requires clarification, as most chemical inhibitors used in clinical trials have dual activity against BACE1 and BACE2 ^35, 39^.

The increased ratios of 1-20&1-34 (BACE2-AβDP) to the amyloidogenic and α-site products are among our most consistent and robust observations in T21 organoid CM and DS-CSF (Fig. 1b-c). The 1-34 generating cleavage can only occur after the cuts by both β- and γ-secretases have released Aβ, because the hidden transmembrane site between aa34 and aa35 is inaccessible to any proteolytic enzymes until the soluble Aβ (1-37 to 1-42) molecules are released from the membrane^12, 13^. Therefore, the Aβ1-34 species can only be a product of an AβDP activity (a catabolic degradation or clearance of an already made Aβ1-37 to 1-42 peptides). Besides BACE2, the only enzymes with potential to cleave the peptide bond Leu34 – Met35 are BACE1^12, 14^, and extracellular matrix (ECM) metalloproteinases (MMP2 and MMP9)^40^, since no other Aβ degrading enzymes (neither IDE, nor NEP, nor ECE) are known to cleave at this site^41^. BACE1 action is unlikely to cause the increased ratios we observe, as BACE1 can only generate this cut in solution at very high enzyme concentration and after prolonged incubation^12^.

To further corroborate this point, we designed a novel FRET-assay and established the conditions in which BACE2 can efficiently cleave at Aβ34 site (Fig. 2) in 2 hours, conditions under which BACE1 activity at the Aβ34 site was undetectable (Supplementary Information). We also demonstrated that two BACE1 inhibitors (β-Secretase Inhibitor IV - CAS 797035-11-1 (Calbiochem, originally a Merck compound)), and LY2886721 (Eli Lilly compound recently used in clinical trials) both inhibit the AβDP activity of BACE2 *in vitro*, while the γ–secretase inhibitor (DAPT) had no effect.

This suggests that the AβDP activity (cutting the peptide bond Leu34 – Met35) has a different enzymatic preference, conditions, and pH, as compared to the classical β-secretase cleavage that both BACE1 and BACE2 are capable of. As FRET assays cleaving this classical (before Asp1) site are generally used to measure the BACE1 inhibitors’ selectivity for BACE1 or BACE2, our data suggest that the degree of selectivity for any given inhibitor calculated this way, does not necessarily reflect whether the same selectivity would apply for their cross-inhibition of the Leu34 – Met35 site cleavage (AβDP) activity. Interestingly, the presence of the Aβx-34 degradation product, both alone^27^ and co-localising with BACE2 (Fig. 4) show elevated levels in cells and extracellular aggregates immediately surrounding neuritic plaques, suggesting BACE2 degradation of not only newly produced Aβ, but also of Aβ that is released and re-deposited (from and to) existing deposits. A recent report on widespread somatic changes in individual neurons suggests an additional mechanism for the production of toxic Aβ species, including products that do not require secretase cleavage^42^, underscoring the importance of efficient Aβ degrading mechanisms that protect from AD, such as the one exerted by BACE2 that we describe here.

A recent mouse model has shown that introducing a third dose of chromosome 21 to a mouse that several hundred fold over-expresses Aβ40 and 42 worsens the amyloid plaque load, and this correlates with an unexpected decrease in the Aβ40/42 ratio^43^. This unfavourable ratio effect (the cause of which is unknown) is expected to worsen the plaque load and AD pathology, and a mere 1.5x increase of *Bace2* dose in this mouse model has no chance in protecting the mouse against a >100x overload of Aβ. In another mouse model, where transgenic *BACE2* was artificially over-expressed together with transgenic wt*APP*, it actually decreased Aβ40 and 42 to the wt mouse control levels, and the presence of *BACE2* transgene reversed behavioural pathologies seen in Tg*APP* mouse^44^.

This indicates that a balance of doses of *APP* and *BACE2* affects levels of soluble Aβ40 and 42, and their oligomerization and aggregation as a consequence. Our results in Figs 5, 6 and 7 further corroborate that a significant disturbance of this balance by a reduction in *BACE2* copy number is sufficient to cause an early AD-like pathology in T21 cerebral organoids. We did not see any amyloid plaque-like structures at >100DIV organoids from three independent T21 iPSC lines (or normal disomic lines) of our isogenic system (Supplementary Figs. 1, 2, 4, 8, 11, 14, Figs. 3, 7, 8). Surprisingly, CRISPR/Cas9 elimination of the third copy of *BACE2* in the same T21 line caused widespread AmyloGlo+ deposits at 41DIV, and widespread neuritic plaque-like structures with profound neuron loss (Supplementary Fig. 11, 12) and Tau pathology at 96DIV (Figs. 6, 7). Our data in Figs.1 and Supplementary Fig. 7 suggest that anti-amyloidogenic activity of BACE2 is gene-dose dependent, and its level varies between individuals, with SNP allelic differences in *BACE2* correlating with age of dementia onset. We therefore hypothesized that organoids grown from some people with DS may develop AD-like pathology without any CRISPR-Cas9 intervention. Diffuse amyloid plaque-like appearance with Tau pathology was recently reported in 110 days old cerebral organoids from only a single DS-hiPSC line^45^ so far. We subsequently analysed iPSC-derived organoids at approximately the same cell culture age from a total of 7 different individuals with DS and one with Dup*APP*. We found flagrant AD-like pathological changes in 5/7 DS tested (71%), as well as the one Dup*APP*. Very interestingly, when this assessment was repeated in independent experiments, and when individual organoids from a single experiment were compared, it was a black/white picture: either they all had AD-like pathology, or none did, driven solely by the genotype of the donor (Table 1). Our data, though not conclusive, are illustrative of the stratifying potential of this technology. For example, the cerebral organoids from individual QM-DS3 showed the worst AD-like pathology with fibrillary amyloid deposits (Fig. 8f, i, j, Table 1), and this individual was diagnosed with dementia at age 37. In contrast, organoids from individual QM-DS6 showed no pathology (Fig. 8b, Table 1), and this individual was also dementia-free at age 37. This opens up possibilities for finding correlations with clinical parameters, for which a much larger number of individuals would have to be tested.

To confirm that the AmyloGlo deposits were in fact aggregated β-amyloid containing material, early organoids were treated with a combination of β-secretase inhibitor IV (βI-IV) and gamma secretase inhibitor XII (Compound E) (Fig. 6 a, b). The combination of these inhibitors should prevent any production of Aβ, and therefore eliminate AmyloGlo positivity. After treatment for 21 days, the inhibitor treatment did indeed prevent the formation of plaque-like deposits within T21C5Δ7 organoids, confirming that such deposits are comprised of β-amyloid. The same treatment conditions also significantly reduced the number of TG3-positive cells in T21C5Δ7 organoids (Fig. 6c), highlighting the ability to modulate both amyloid and tau pathology in the cerebral organoid system. This also demonstrates the feasibility of using this AD-like organoid pathology in future hypothesis-free drug screens for chemical compounds that may prevent/inhibit amyloid production or aggregation.

In view of our results, it becomes inviting to hypothesize that triplication of *BACE2* may be the cause of the delayed onset of dementia in 30% of people with DS compared to Dup*APP*^7^, and (because of the predicted abundance of BACE2 mRNA in endothelial cells) also the cause of a significantly lower degree of cerebral amyloid angiopathy (CAA) in the brains of people with DS compared to those of Dup*APP*^46^. Our organoid system is not informative in this regard, as we could not detect any endothelial cells in our organoids (not shown). This, however, is also an advantage, as it allows uncovering the mechanisms that are specific to neurons in the absence of endothelial or blood cell derived tissue components.

In neurons, a recent report also found that an increased *APP* dose may act (through an unknown mechanism) as a transcriptional repressor of several chromosome 21 genes, including *BACE2*^47^. This observation needs further verification and mechanistic explanation, but if true, it would imply that the protective effect of the third copy of *BACE2* in DS that we observe is actually quenched by the third copy of *APP*, which opens up possibilities of chemically intervening to inhibit this transcriptional repression and potentially unleash a much greater degree of BACE2 protection. An integration of the two observations (the one in^47^ and the one in our report) suggests this could be exploited as an additional new protective/therapeutic strategy for AD in general.

We found, surprisingly, an equally high or higher level of colocalisation of Aβx-34 with LAMP2A, as with the general LAMP2 (Fig. 3). The high level of colocalisation with LAMP2A and absence of colocalisation with either LC3A or LAMP1 (Fig. 3), suggest that AβDP activity of BACE2 that generates Aβ34 is not related to classical lysosomal degradation or macroautophagy, but rather could be related to a CMA-like process^48, 49^. The only published study that linked CMA with APP processing^50^ found a motif that satisfies the criteria for a CMA-recognition KFERQ motif at the very C-terminus of APP (KFFEQ), and this paper demonstrated that C99 (β-CTF) can bind HSC70. However, paradoxically, when this motif is deleted from the β-CTF, the binding to HSC70 is not abolished, but rather increased, suggesting the presence of another, alternative CMA-recognition motif within the β-CTF peptide^50^. The association of the AβDP x-34 product with LAMP2A/CMA compartment is a provocative new observation that requires further studies.

In conclusion, we found that relative levels of specific non-amyloidogenic and AβDP (Aβ-clearance) products are higher in T21 organoids and DS-CSF, and they respond to the dose of *BACE2* (and not *APP*). We also demonstrated that BACE2-AβDP activity generating one of these products can be cross-inhibited in solution by recently clinically tested BACE1-inhibitors. All components of the AβDP degradation reaction (hitherto only demonstrated in solution *in vitro*): the main substrate (Aβx-40), the enzyme (BACE2), and its putative degradation product (Aβx-34), we found highly colocalised in discrete intracellular vesicles in human brain neurons, (and not astrocytes), suggesting that at least some of the AβDP activity generating Aβx-34 takes place intra-neuronally and physiologically during lifetime, before the onset of AD pathology, in both normal and DS brains. Furthermore, we directly demonstrated that the third copy of *BACE2* protected T21-hiPSC organoids from early AD-like amyloid plaque pathology, therefore proving the physiological role of BACE2 as an AD-suppressor gene. The BACE2’s θ-secretase anti-amyloidogenic cleavage and the AβDP degradation actions could both be contributing to an overall AD-suppressive effect. Regardless of the contribution of each of these modes of action, our combined data suggest that increasing the action of BACE2 could be exploited as a therapeutic/protective strategy to delay the onset of AD, whereas cross-inhibition of BACE2-AβDP activity by BACE1-inhibitors would have the unwanted worsening effects on disease progression. We also show that cerebral organoids from genome-unedited iPSCs could be explored as a system for pre-morbid detection of high-risk population for AD, as well as for identification of natural dose-sensitive AD-suppressor genes.

## Methods

### Human subjects – clinical assessment

Human subjects were participants in the “The London Down Syndrome Consortium (LonDownS): an integrated study of cognition and risk for Alzheimer’s Disease in Down Syndrome” inter-disciplinary study, enrolled after informed consent, as per ethical approval 13/WA/0194, IRAS project ID:120344 see http://www.ucl.ac.uk/london-down-syndrome-consortium). Dementia diagnostic status was obtained via carer report and medical records, based on assessment by the individual’s own clinician. To confirm diagnoses, we collected detailed data on dementia symptoms from the Cambridge Examination for Mental Disorders of Older People with Down’s Syndrome and Others with Intellectual Disabilities (CAMDEX), a clinical tool used for diagnosing dementia in individuals with DS. This was independently reviewed by two psychiatrists. One iPSC line (QM-DS1) was established from an individual 40 year old who was diagnosed with dementia aged 40. Consensus ratings agreed that this individual showed signs of dementia and died two years later following significant decline. AD was noted as the cause of death. Another line came from an individual (QM-DS2) who was diagnosed with dementia aged 63. The sample was taken at age 67 and at this time point, consensus was that the individual presented with signs indicative of possible dementia. This individual was still alive for follow-up two years later, with a confirmed diagnosis of AD-dementia and AD-related seizures developed at age 68. The third line came from an individual diagnosed with dementia at age 37 (QM-DS3). Another line came from the individual QM-DS6, who donated hair at age 31, and remains dementia free after follow up at age 37. The age, sex, ApoE genotype and dementia status of all other individuals are detailed in Table 1.

### Primary hair follicle keratinocyte sampling

Upon specific informed consent, three to six individual strands of hair were non-invasively plucked from the scalp hair of donor subjects, and placed in transport medium [DMEM (Sigma D5546), 2mM glutamine (Sigma G7513), 1x Pen/Strep (Sigma, P4333), 10% foetal calf serum]. Upon arrival to the lab, hair follicles were placed in collagen coated T25 flasks in KGM2 medium (Lonza CC-3107) and incubated at 37°C, 5% CO2. Primary keratinocyte cultures were split after reaching 35-50% confluency using 0.05% Trypsin/0.02% EDTA.

### Reprogramming of primary keratinocytes

Primary keratinocyte cultures were expanded to 70% confluency, electroporated with plasmids encoding reprogramming factors in episomal vectors (non-integrational reprogramming), and transferred to 0.1% gelatin coated wells (6 well plate), pre-seeded with mitotically disabled mouse embryonic fibroblast (MEF) feeder cells. Specifically, trypsinised 700,000 keratinocytes were washed once with sterile PBS and electroporated using the Nucleofector 4D (Lonza, X-module apparatus, kit V4XP-3024, programme DS138, following manufacturer’s instructions) with 3 µg of the episomal plasmid mix (equimolar mixture of plasmids obtained from Addgene: pCE-hOct3/4, pCE-hSK, pCE-hUL, pCE-mp53DD and pCXB-EBNA1). After electroporation, cells were transferred from the cuvette to KGM2 medium (Lonza CC-3103 and CC-4125).

Solution was gently mixed and transferred to the 6 well plate with feeders. On day 2 (48 hours after electroporation) medium was removed and replaced with fresh KGM2 medium. On the day 4, the medium was switched to standard human embryonic stem cell (hESC) medium (High glucose DMEM with 20% Knockout Serum Replacement, non-essential amino acids, Glutamax+ penstrep (Life Technologies), 2-mercaptoethanol (100µM) with 10-20 ng/mL of FGF2). From day 10 onwards, MEF-conditioned medium supplemented with 10-20 ng/mL of FGF2 was used. By day 20, iPSC colonies were observed. After day 30, large iPSC colonies were mechanically picked, and expanded using ReLeSR and hESC medium plus ROCK inhibitor (Y-27632, Stem Cell Technologies). The iPSC lines were generated in this way from six unrelated people with DS (QM-DS1, QM-DS2, QM-DS3, QM-DS4, QM-DS5 and QM-DS6 respectively, detailed in Supplementary Table1). The iPSCs from a 64 year old euploid patient with FEOAD caused by Dup*APP*^20^ (QM-Dup*APP*) were generated from peripheral blood mononuclear cells, using the same non-integrational episomal reprogramming vectors as above, and a modified protocol.

Specifically, 10^6^ PBMCs were electroporated with 3ug of episomal plasmids (equimolar) using program EO-115 and solution p3 on the Amaxa 4D nucleofector. Electroporated PBMCs were transferred into 1 well of a 6 well plate seeded with MEFs in PBMC recovery medium (RPMI supplemented with 200uM 1-thioglycerol (Sigma M6145), 1uM Dexamethasone (Sigma D1756), 2U/ml Erythropoietin (R&D Systems 287-TC-500), 100ug/ml Holo-transferrin (R&D Systems 2914-HT), 40ng/ml IGF-1 (Peprotech 100-11), 10ng/ml IL-3 (Peprotech 200-03), 100ng/ml SCF (Peprotech 300-07). After 2 days, 2ml of PBMC recovery medium was added to transfected cells. After another 2 days, PBMC recovery medium was replaced every second day with hESC medium supplemented with 10ng/ml FGF-2. Visible iPSC colonies were mechanically picked and expanded as per the keratinocyte reprogramming protocol. The iPSCs for the isogenic DS model included isogenic lines derived from an individual with Down Syndrome mosaic for T21 were described by us previously^17^: D21-C3, D21-C7, D21-C9 and T21-C5, T21-C6, T21-C13. These were generated by non-integrationally reprogramming primary skin fibroblasts from a person with mosaic DS, using Sendai virus delivered standard Yamanaka OKSM factors. The total number of iPSC lines generated per patient is listed in Supplementary Table 1. iPSCs were maintained on Geltrex coated plates and cultured in E8 medium (Life Technologies) supplemented with penicillin/streptomycin. Passaging was carried out using ReLESR and 10 μM ROCK inhibitor was included in culture media for 24 hours after passaging.

### Cerebral organoids

Cerebral organoids were generated following the standard protocol with the following changes^18^. iPSC lines were first transitioned into feeder free conditions using either mTESR1 or E8 media with geltrex. To form embryoid bodies (EBs), hiPSCs were washed once with PBS, then incubated with Gentle Cell Dissociation Solution (Stemcell Technologies) for 4mins. This solution was then removed and accutase added and incubated for a further 4mins. mTESR1/E8 medium at double the volume of accutase was added to the cells and a single cell suspension generated by titruating. Cells were centrifuged to remove accutase and then resuspended in hESC medium supplemented with 4ng/ml FGF2 and 50micromolar ROCK inhibitor. 9000 cells were used to form a single EB in each well using either a V shaped ultra low attachment 96 well plate (Corning). Specifically, iPSCs were allowed to form embryoid bodies (EBs) in suspension by culturing for 6 days in hESC medium with low FGF, in non-adherent culture dishes. After 5-7 days, EBs were transferred into a 24 well ultra low attachment plate for neural induction. Neural induction was achieved by culturing for further 5-7 days in DMEM-F12 supplemented with 1% of each: N2, GlutaMAX and MEM-NEAA, plus 1μg/ml heparin.

Neurally induced EBs showing neuroectodermal “clearing” in brightlight microscopy were embedded in matrigel droplets, and transferred to 6cm dishes containing organoid differentiation medium-A, (for 4-5 days), followed by organoid differentiation medium+A^18^. Organoid maturation was carried out with 12-16 organoids per 6cm dish on an orbital shaker at 37°C, 5% CO_2_. Aliquots of conditioned medium (CM) were collected from mature organoids (100-137 days old from day of EB formation), 3-4 days after feeding (to allow time for cells to secrete products into the culture media). Three completely independent experiments were carried out each time starting from undifferentiated iPSC stage, and CM was collected at 3-4 timepoints in each experiment. CM was immediately frozen and stored at −80°C.

For inhibitor treatment, organoids were treated from 20DIV (6 days after embedding in matrigel) to 41DIV. βI-IV and Compound E were added freshly to the media before use at final concentrations of 2.5μM and 6nM respectively. Media was replaced every 3-4 days during treatment. DMSO of the same volume was used as a vehicle only control.

### IP-MS

CM from organoids was analysed by IP-MS, using a previously described method^20^. The team performing the MS was blinded to the genotypes in all experiments. In exp1, all three independent trisomic lines (T21C6, T21C5, and T21C13) were compared to two independent disomic lines (D21C3 and D21C7), whereas in exp2, two independent trisomic lines (T21C6 and T21C13) were compared to two independent disomic lines (D21C7 and D21C9).

In exp3, a T21C6 line was compared to the isogenic D21C9 line, and to hiPSC lines from 3 unrelated individuals: a Dup*APP* FEOAD patient (QM-Dup*APP*), and two unrelated adult people with DS (QM-DS1 and QM-DS2).

In all 3 experiments, IP-MS results for all iPSC lines that were used in a particular experiment are shown. IP-MS results were used to calculate the relative ratios of peptides and these ratios were taken as data points for the statistical comparisons.

IP-MS spectra were also obtained from the CSF samples of people with DS and age-matched normal controls. Peak ratios calculated as described above. The cohorts, methods and spectra behind these data were previously described^26^.

### Fluorescence in situ Hybridization (FISH)

FISH on organoid cryosections was performed as described^51^. Briefly, slides were rinsed in PBS, rehydrated in 10 mM sodium citrate buffer and incubated in the same buffer at 80⁰C for 20 min. Slides were cooled down and incubated in 2x Saline Sodium Citrate (SSC) for 5 min and in 50% formamide in 2x SSC for 1h. After incubation slides were covered with previously prepared hybridization chamber and incubated with 10 μL of XA 13/18/21 (D-5607-100-TC, MetaSystems Probes) Probe, protected from the light, at 45⁰C for 2h, at 80⁰C for 5 min and for 2 days at 37⁰C in the water bath. Slides were rinsed with 2x SSC at 37⁰C (3×15 min) and with 0.1x SSC at 60⁰C (2×5 min) on the shaker, equilibrated in 2x SSC at 37⁰C for 2 min, counter-stained with DAPI for 10 min and covered with Dako Fluorescent Mounting Medium. Fluorescence was captured on Zeiss LSM-800 inverted confocal microscope with Airyscan using 63x oil-immersed objective. Image analysis was performed using IMARIS x64-v9.1.2. Software (BITPLANE, An Oxford Instruments Co., Zurich, Switzerland).

Quantification was performed on 10-13 figures, with >1500 cells per genotype. The spots specific for chromosome 21 were labelled in the red spectrum, whereas the spots specific for chromosome 13 were labelled in the green spectrum. More than 500 nuclei from eighth different Z-stacks were evaluated for each line. Based on the observed number of fluorescent hybridization signals, nuclei were assigned to four different categories, namely ‘one signal’, ‘two signals’ ‘three signals’ and ‘> three signals’. Damaged nuclei or overleaped nuclei with other nuclei were not included in scoring.

### Immunostaining of organoids

Cerebral organoids derived from iPSCs and grown for indicated number of days *in vitro* (DIV) were fixed in 4% PFA, cryoprotected in 30% sucrose/PBS solution and embedded in OCT. Twenty micron thick sections were cut and mounted on Superfrost Plus slides (Fisher Scientific) for immunostaining.

For immunofluorescent staining, permeabilisation and blocking was carried out in 3% donkey serum with 0.2% TritonX-100 in PBS for 1h at room temperature (RT). Primary antibodies (Supplementary Table 3) were diluted in 1% donkey serum with 0.2% TritonX-100 in PBS and incubated overnight at 4°C. Following washes with PBS, secondary antibodies (Supplementary Table 4) were diluted in 1% donkey serum with 0.2% TritonX-100 in PBS and incubated for 2h at RT. Following washes with PBS, sections were counterstained with DAPI or DR (Supplementary Table 2) for 10 min at RT, washed again and mounted with DAKO fluorescent mounting medium. As negative controls for all antibodies, secondary antibody only controls were carried out (Supplementary Fig. 14).

### Western Blot

For western blots, whole cell lysates of CRISPR edited or unedited iPSCs (Fig. 5c) or organoids (Fig. 7o) were separated in a 10% acrylamide gel by SDS-PAGE and transferred to a nitrocellulose membrane according to the manufacturers protocols (Bio-Rad). Following a 60min incubation in 5% non-fat milk in TBS-T the membrane was incubated with primary and secondary antibodies (Supplementary Tables 3, 4). For the stainings shown in Fig. 7o quantitations were done strictly on the same membranes re-stained using the antibodies shown. For the protein of interest (BACE2 or TG3), the signal was adjusted to corresponding β-actin loading control for all samples. Such adjusted values for unedited C5 (wt) (n=4) were set to 1, and used to calculate the fold change for C5Δ7 (n=4) replicates, and the resulting fold-change values for pairs run on the same gel were averaged and analysed by student’s t-test. Membrane stripping between stainings was carried out using Thermo-Fisher stripping solution, following manufacturer’s instructions.

### AmyloGlo and Thioflavine S staining

For AmyloGlo staining, OCT embedded slices were rinsed with PBS, and incubated in 70% ethanol for 5 min at RT, followed by washing with milli Q water for 2 min at RT. Slices were then incubated with AmyloGlo solution for 10 min in the dark at RT, followed by washing in 0.9% saline solution for 5 min at RT, and counterstaining with DR for 10min at RT. Thioflavine S staining was performed as described^52^. OCT embedded slices were rinsed with PBS and incubated with Thioflavine S solution for 10 min in the dark at RT, differentiated in 80% ethanol and counterstained with DR for 10 min at RT, rinsed with PBS and mounted with DAKO fluorescent mounting medium.

### Formic acid pre-treatment

To increase signal of insoluble β-amyloid material T21C5Δ7 (96DIV) organoid slices were treated with 87% Formic acid for 10 min at RT. After 10 min Formic acid were rinsed three times with PBS and samples were immunostained as described above.

### Human brain samples

PFA-fixed, paraffin-embedded human anonymized post-mortem brain samples were obtained from the Brain Bank of the Croatian Institute for Brain Research (CIBR), Institute of Pathology of The Royal London Hospital (IP-RLH) and the South West Dementia Brain Bank (UK) (Supplementary Table 1). Slices were cut at 5-10 µm thickness, and stained using primary antibodies, secondary fluorophore-coupled anti-Ig antibodies and their respective dilutions (Supplementary Tables 3, 4).

### Immunohistochemistry and immunofluorescence on human brain tissue

PFA-fixed, paraffin-embedded 5-10 µm thick slides were de-paraffinised by incubation in xylene, rehydrated in a graded series of ethanol and rinsed in PBS for 10 min. For antigen retrieval, the slides were steamed in 0.01M citrate buffer, pH 6.0 at 100⁰C for 30 min, cooled and rinsed 3×10 min in PBS. On slides used for IHC, endogenous peroxidases were quenched with 0.025% hydrogen peroxide for 30 min at RT and rinsed 3×10 min in PBS. Double immunohistochemical staining was performed “Polink DS-MR-Hu A2 Kit for Immunohistochemistry Staining” (GBI Labs, DS202A-18). Briefly, Polymer-HRP and AP Double staining kit distinctly labels two different antigens in human tissue, using mouse (GBI-Permanent Red) and rabbit (DAB-brown) antibodies. Single IHC was performed “VECTASTAIN ABC HRP Kit”. For immunofluorescence, following antigen retrieval, slides were incubated in blocking/permeabilisation solution (0.2% Triton X-100 in PBS + 3% donkey serum) for 1h at RT. The slides were incubated over night at 4⁰C with primary antibodies (Supplementary Table 3) in solution (0.2% Triton X-100 in PBS + 1% donkey serum). Next day primary antibodies were rinsed 3×5 min in PBS and incubated for 2h with secondary antibodies (Supplementary Table 4) in 0.2% Triton X-100 in PBS at RT and rinsed 3×5 min in PBS. Nuclei were counterstained with DAPI for 10 min, rinsed 3×5 min in PBS and mounted with Dako Fluorescent Mounting Medium. In order to distinguish the contribution of lipofuscin auto-fluorescence to the colocalised signals, specificity of primary antibodies (Aβx-34 and BACE2) has been validated using three different methods: Sudan black B staining (Supplementary Fig. 15a), pre-incubation with BACE2 specific immunogenic peptide (Supplementary Fig. 15b-e) and Lambda (λ) scan function on confocal microscope (Supplementary Fig. 15f, g). Three different samples (DS-AD1, DS (28 yrs) pre-AD and euploid sporadic AD (73 yrs) after IHC were stained with 0.1% Sudan black B in 70% ethanol for 20 min at RT and analysed on confocal microscope with Aiyrscan. Sample DS-AD1 was stained with antibodies solution, 12 hrs pre-absorbed with BACE2 specific immunogenic peptide, and analysed on confocal microscope and slide scanner. Lambda scan records a series of individual images within a defined wavelength range (in our case from 630 nm to end of spectrum) and each image was detected at a specific emission wavelength, at 10 nm intervals. For lambda scan analysis, samples were stained with one primary antibody and labelled with far-red secondary antibody (647). As negative control, we used secondary antibody (647) alone and, as additional negative control, one sample was counterstained with DAPI only, without secondary antibody. As we used a far-red (647) antibody, we analysed expression from 630 nm to the end of spectrum at 10 nm intervals. Aβx-34 and BACE2 antibodies showed specific peaks, significantly over and above the autofluorescent signal, in all three specific ROI indicated in Fig. 4 and Supplementary Fig. 15 (intraneuronal fine-vesicles, large intraneuronal spherical granules and extracellular aggregates). DAPI and secondary antibody alone show peaks only at background level. Samples were analysed by: LSM800 Inverted Confocal Microscope with Airyscan (ZEISS), LSM800 Upright Confocal Microscope (ZEISS), LEICA DM6000 CFS and Axioscan.Z1 Slide Scanner (ZEISS). As negative controls for all antibodies, secondary antibody only controls were carried out (Supplementary Fig. 15).

### Gallyas staining

For Gallyas staining samples were depariffinised and/or rinsed in PBS, then treated with Ammonium-Silver Nitrate (0.1 g NH_4_NO_3_, 0.1g AgNO_3_, 0.3 mL 4% NaOH) solution for 30 min protected from the light, rinsed with 0.5% acetic acid (3 x 3 min) and placed in developer solution for 5-30 min. Developer solution was made from three stock solutions: 25 ml of Solution A (50g Na_2_CO_3_ + 1000 mL distilled water), 7.5 ml of Solution B (2g NH_4_NO_3_ + 2g AgNO_3_ + 10g Tungstosalicic acid hydrate + 1000 mL distilled water) and 17.5 ml of Solution C (2g NH_4_NO_3_ + 2g AgNO_3_ + 10g Tungstosalicic acid hydrate + 7.3 ml 37% formaldehyde solution +1000 mL distilled water). After developer solution samples were rinsed in water and placed in destaining solution (30g K_2_CO_3_ + 55g EDTA-Na_2_ + 25g FeCl_3_ + 120g Na_2_S_2_O_3_ + 20g KBr +1000 mL distilled water). Finally, samples were rinsed two times in 0.5% acetic acid. After staining samples were rinsed in water, dehydrated in a graded series of ethanol, cleared in Histo-Clear and mounted with Histomount mounting medium. Samples were scanned by NanoZoomer 2.0RS (HAMAMATSU).

### Image Analysis

Immunofluorescent stains of 20µm thick slices are shown as maximal projections captured on Zeiss LSM-800 upright confocal microscope using 63x oil-immersed objective. Image analysis was performed using IMARIS x64-v9.3.1. Software (BITPLANE, An Oxford Instruments Co., Zurich, Switzerland). Quantification was performed blinded to the genotype, on 5 independent images representing 3 individual organoids per genotype, and containing 3,000-4,000 cells per image. Only images within the “cortical” part of the organoid were considered for analysis. For quantification of protein/peptide markers, total fluorescence intensity of positive signals for each wavelength for a given antibody was normalised to the total fluorescence intensity for MAP2 as a pan-neuronal marker.

For colocalisation calculations: Image analysis was performed using IMARIS software. Pairwise Pearson’s coefficient of colocalised volume for a pair of costained antibodies with contrasting fluorescence wavelengths was automatically calculated by the IMARIS software on 3-8 images from 3 independent organoids, per any given antibody combination, using a maximal projection through the entire z-stack.

### A new FRET-assay for the detection of AβDP activity generating Aβ1-34

We designed and synthesized a new FRET-based peptide containing the fluorophore at one end and the quencher at the other end, and spanning the Aβ34 site in its middle. The exact peptide design is under discussion for intellectual property protection. The FRET (BACE2 R&D Systems, not based on amyloid sequence) control peptide (10 µM) (not shown) or the newly designed FRET Aβ34-site peptide (10 µM) were digested at 37° C by human BACE2 (R&D systems, 1 ng/µL) in presence or absence of the stated inhibitors for 2h. Enzyme activity was defined by measuring the fluorescence increase before and after the incubation. After blank-subtracted fluorescence units where normalized to the control digest, one-way ANOVA was performed. P-values were calculated with a *post-hoc* Bonferroni correction for multiple comparisons (only pairs relative to the untreated control were simultaneously compared). Error bars represent standard error.

### Statistical Analysis

Initial analysis was carried out using Microsoft Excel to calculate two-tailed student t-tests. Additional Holm-Bonferroni correction was carried out using the Excel macro from ref^53^. For all multiple comparison analysis, ANOVA and Holm-Bonferroni calculations were performed at http://astatsa.com/OneWay_Anova_with_TukeyHSD/

### SNP arrays

iPSC lines genome integrity: Genomic DNA was isolated from iPSCs using standard column kits. DNA of all iPSC lines shown in the manuscript were re-analysed at the similar passage used for the derivation of organoids using SNP arrays Illumina OmniExpress v1.1 chips and analysis performed in Genome Studio 2.0 software. Following CRISPR editing, C5Δ7 was assessed by SNP array and no genomic alternations were detected compared to the parental C5 iPSC line. See the raw data from this for chromosome 21 array, the rest are not shown for the lack of space, data available on request). The genome integrity of the isogenic iPSC lines was previously published^18^ (but was repeated here as described above). No additional rearrangements due to re-programming or passaging were observed.

*BACE2* locus SNPs: The cohort of people with DS has been described in recent reports ^22, 23^. In brief, participants donated DNA samples and had detailed cognitive and clinical assessments to determine dementia status^54^. Age of dementia diagnosis was established and used in SNP analysis. *BACE2* SNP genotyping for the LonDownS cohort was undertaken as previously described^24^. Briefly, 93 single nucleotide polymorphisms (SNPs) located within the *BACE2* locus +/-50kbp, were genotyped in 554 individuals recruited through the LonDownS Consortium. Genotyping was done in the UCL Genomics Centre using Human OmniExpressExome v1.2,v1.3,v1.4 beadchips. SNP clustering and genotyping, was undertaken using GenomeStudio (Illumina, San Diego, CA, USA). Manual reclustering for Chr 21 SNPs was done using GenomeStudio module v1.9.4 polyploidy-genotyping (http://res.illumina.com/documents/products/technotes/technote_genomestudio_polyploid_genotyping.pdf). Comparison of SNP MAF for the European subset of the LonDownS cohort versus the reference European unaffected cohort was undertaken to ensure no overrepresentation of SNPs due to the genetic background. Age at onset (or Age of onset (AOO)) regression was analysed using an additive model with dosage of Top A allele of each SNP as independent variable. Sex, APOE4 dosage, and 8 PCA component of genetic variation were used as covariates. A nominal p value of 0.05 was used as a cutoff for nominal significance. Two SNPs in the *BACE2* locus (purple, Supplementary Fig. 7) were nominally associated with AOO in the LonDownS cohort, but were not significant after correction for multiple testing.

### Quantitative paralogous amplification-pyrosequencing

Quantitative paralogous amplification-pyrosequencing was carried out based on the published method^55^. This method takes advantage of the existence of identical sequences on chromosome 21 and one other autosome, allowing amplification of both loci with a single primer pair. Paralogous sequence mismatches in amplified products from chromosome 21 (GABPA and ITSN) can be quantified relative to their paralogous regions on chromosome 7 and 5 respectively. As such, trisomic cells show a 60:40 ratio for the paralogous sequence, while disomic cells produce a 50:50 ratio. Primers used are listed below, and pyrosequencing was performed on the Pyromark Q48 machine (Qiagen) following standard procedures.

### CRISPR/SpCas9-HF1Cas9 editing of the *BACE2* locus

The guide-RNA (gRNA) targeting *BACE2* Exon 3 was cloned into a vector containing the high fidelity SpCas9-HF1^56^ and blasticidin S resistance gene. The complete plasmid was delivered via Lipofectamine3000 to a trisomic iPSC line T21C5 (full official name NIZEDSM1iT21-C5), which was described and characterized in a previous report^18^. Untransfected iPSCs were removed by treatment with blasticidin (2 μg/ml for 48h). Individual colonies were picked and further sub cloned by limiting dilution to achieve clonal cell lines. DNA was purified from individual clones, PCR amplified and sequenced by Sanger Sequencing. Sequences were analysed in Mutation Surveyor (V3.1.0) and “Tracking InDels by dEcomposition (TIDE)” (TIDE V 2.0.1, Desktop Genetics). TIDE analysis of the CRISPR-targeted clone 2.3.5 DNA sequence gave a score of 65% of the wt read remaining (not shown). The quality of the gRNA was assessed using two different prediction software platforms: CCTop online software^57^, and the MIT online platform (http://crispr.mit.edu/). The same two software platforms were used to predict the off-target sites.

Neither platform found any off-targets with 0, 1 or 2 mismatches. The top 10 CCTop-predicted sites were PCR amplified in both Δ7 and WT clones, then sequenced by Sanger Sequencing to rule out off target events. No differences in the sequence were found.

### Protein isolation from Cortical Organoids

Organoids were collected at specified durations in culture (expressed as Days *In Vitro* (DIV)) and washed twice with ice-cold PBS. The samples were resuspended in ice-cold NP-40 Buffer (150mM NaCl, 1% NP-40, 50mM Tris pH8) containing EDTA free protease inhibitors (complete cocktail, Roche) and lysed using a 1ml tissue homogenizer (Fisher). Each sample was centrifuged at 10,000rpm for 10 minutes at 4°C and the homogenates were stored at −80°C. Protein concentration was determined using the bicinchoninic acid method (BSC, Pierce).

### Detection of fibrillary material from organoids by transmission electron microscopy (TEM)

Organoids were lysed following the same procedure for protein extraction, however, samples were initially spun at 20,000g for 20 minutes at 4°C. Following the first centrifugation, supernatants were removed and kept on ice. The remaining cell pellets were resuspended in 5x weight/volume buffer (10mM Tris-HCL pH7.5, 0.8M NaCl and 10% sucrose)^58^ containing proteases inhibitor and spun at 20,000g for 20 minutes at 4°C. An equal volume of supernatant 1 was added to the supernatant from the second centrifugation step. 1% N-lauroysarcosinate (weight/volume) was added and the samples were rocked at room temperature for one hour. The samples were ultra-centrifuged at 100,000g for one hour at 4°C. The supernatant was decanted and the sarkosyl-insoluble pellet was resuspended in ice cold PBS prior to imaging. The samples were deposited on to glow-discharged 400 mesh formvar/carbon film-coated copper grids.

Negatively stained with a 2% aqueous (w/v) uranyl acetate solution and then immediately analysed at 100 kV using a JEOL TEM1010 equipped with a Gatan Orius camera.

### TEM analysis of synthetic Aβ1-40 fibrils *in vitro*

Synthetic Aβ peptide powder (China peptides) was treated with 1,1,1,3,3,3-hexafluoro-2-propanol (HFIP) and lyophilized. The peptide was then dissolved in 20µL of 100 mM NaOH and then diluted with buffer. A 50µM stock of this monomeric Aβ peptide was grown at 37°C shaking at 180 rpm for 48-60 hours before recording the TEM images. 4µL of extract was added to a 15 nm thick, lacey carbon on 300 mesh grid (glow-discharged) for 2 minutes followed by negative staining with 2% uranyl acetate for 1 minute and then air dried. The grids were then viewed under FEI T12, 120 kV Transmission electron microscope equipped with a 4K CCD camera (FEI) at 30000X magnification under low dose conditions.

## Appendix 1

LonDownS Consortium, The Wellcome Trust, London, UK. The LonDownS Consortium principal investigators are: Andre Strydom, Department of Forensic and Neurodevelopmental Sciences, Institute of Psychiatry, Psychology and Neuroscience, King’s College London, London, UK and Division of Psychiatry, University College London, London, UK; Elizabeth Fisher and Frances Wiseman, Department of Neurodegenerative Disease, UCL Institute of Neurology, London, UK; Dean Nizetic, Blizard Institute, Barts and the London School of Medicine, Queen Mary, University of London, London, UK, and Lee Kong Chian School of Medicine, Nanyang Technological University, Singapore, Singapore; John Hardy, Reta Lila Weston Institute, Institute of Neurology, University College London, London, UK, and UK Dementia Research Institute at UCL, London, UK; Victor Tybulewicz, Francis Crick Institute, London, UK and Department of Medicine, Imperial College, London, UK; and Annette Karmiloff-Smith (Birkbeck University of London) (deceased).

## Data availability

All data that support the findings described in this study are available within the manuscript and the related supplementary information, and from the corresponding authors upon reasonable request.

## Acknowledgments

DN’s work was funded by the Singapore National Medical Research Council (NMRC/CIRG/1438/ 2015), Singapore Ministry of Education Academic Research Fund Tier 2 grants (2015-T2-1-023 & 2015-T2-2-119), and EU-JPND - “Heroes” and “CoEN” Consortia. DN, JH and AS all received funding as part of The Wellcome Trust “LonDownS Consortium” Strategic Funding Award (098330/Z/12/Z) (UK), and JH received funding from the Dementia Research Institute, an anonymous foundation and the Dolby foundation. HZ is a Wallenberg Academy Fellow supported by grants from the Swedish Research Council, the European Research Council, Swedish State Support for Clinical Research (ALFGBG-720931) the UK Dementia Research Institute at UCL. KB holds the Torsten Söderberg Professorship in Medicine at the Royal Swedish Academy of Sciences, and is supported by the Swedish Research Council (#2017-00915), the Swedish Alzheimer Foundation (#AF-742881), Hjärnfonden, Sweden (#FO2017-0243), and the Swedish State Support for Clinical Research (#ALFGBG-715986). AM was awarded a William Harvey Academy Fellowship, co-funded by the People Programme (Marie Curie Actions) of the European Union’s Seventh Framework Programme (FP7/2007-2013) under REA grant agreement n° 608765. JNF received a fellowship from the Singapore National Research Foundation (NRF-NRFF2016-03). ARL received funding from the Fondation pour la Recherche Médicale (FRM). JGh and AR are supported by the BrightFocus Foundation (A2015275S) and NIH grant AG059695. The work of ŽK, GŠ, IK and DM research was co-financed by the Scientific Centre of Excellence for Basic, Clinical and Translational Neuroscience (project “Experimental and clinical research of hypoxic-ischemic damage in perinatal and adult brain”; GA KK01.1.1.01.0007 funded by the European Union through the European Regional Development Fund). ŽK is also supported by the Adris Foundation. GŠ is also supported by the Croatian Science Foundation (HRZZ IP-2019-04-3584). DM is supported by the Croatian Science Foundation (IP-2016-06-9451). The South West Dementia Brain Bank is jointly funded by Alzheimer’s Research UK and Alzheimer’s Society, and is supported by BRACE (Bristol Research into Alzheimer’s and Care of the Elderly) and the Medical Research Council, UK. All unique materials will be made available for academic and non-commercial research purposes. We acknowledge Balakrishnan Kannan in the LKC imaging facility for his assistance. The authors thank Géraldine Joly-Hélas, Pascal Chambon, Željka Punčec, Ana Bosak and Danica Budinščak for technical help, Marie Loh and Jacqueline Tai for the use of pyrosequencer, Moses Tandiono for help with SNP-array experiments, Selina Wray for some antibodies, and Anna Barron and Madeline Lancaster for advice. We are grateful to Maria Grazia Spillantini for advice, critical comments, and AT100 antibody.

## Author Contributions

IA, PAG, AM, EP, JH, HZ and DN contributed to the concept and design, the acquisition, analysis and interpretation and drafting of manuscript

AS, DK, JGr, SHav, NRD and KB contributed to the concept and design and to acquisition and analysis and interpretation

YJY, GB, RH, CS, SHam, DW, ARL, HK, JEM, AR, JGh, ŽK, GŠ, and LC contributed to acquisition of data

GG, KYM, RB, NLO, MP, KP, MT, AS, DLB and DM contributed to acquisition, analysis and interpretation of data

EG, XS, HK and JNF contributed to acquisition and analysis of data

EV contributed to analysis and interpretation

IK and PTF contributed to the concept and design and the acquisition of data

## Supplementary Figures

**Supplementary Fig. 1.**
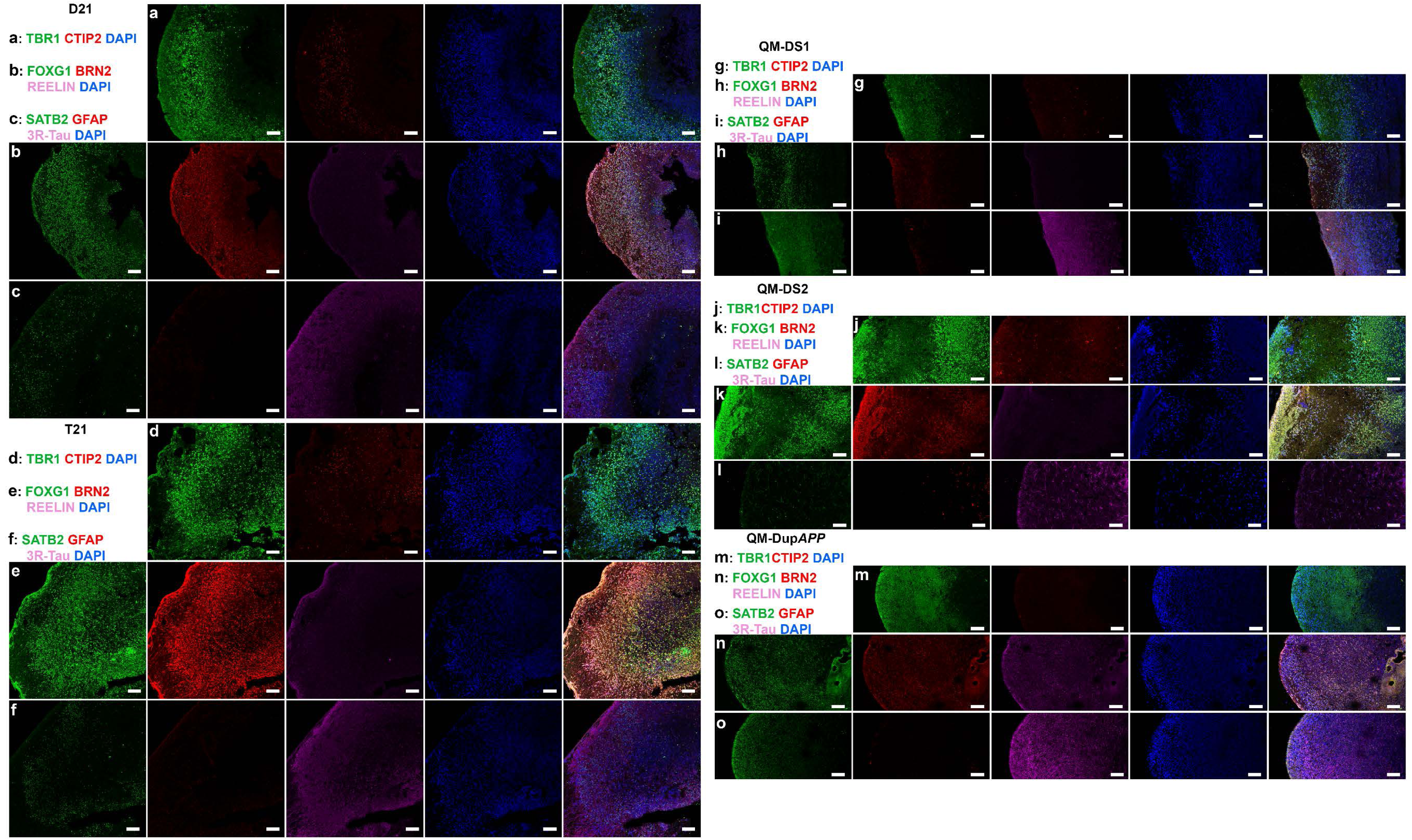
Cerebral organoids express cortical neuronal layer-specific and astrocyte markers. Representative images of isogenic D21 (**a-c**), isogenic T21 (**d-f**), and non-isogenic QM-DS1 (T21) (**g-i**), QM-DS2 (T21) **(j-l**) and QM-DupAPP (**m-o**) organoids are shown, confirming expression of TBR1 (layer IV), CTIP2 (layer V), FOXG1 (layers II and III), BRN2 (layer VI), REELIN (layer I), SATB2 (layer III), GFAP (astrocytes) and 3R-Tau (neurons). Final image in each row is a merge of individual antibodies in that row. Scale bar: 100μm.

**Supplementary Fig. 2.**
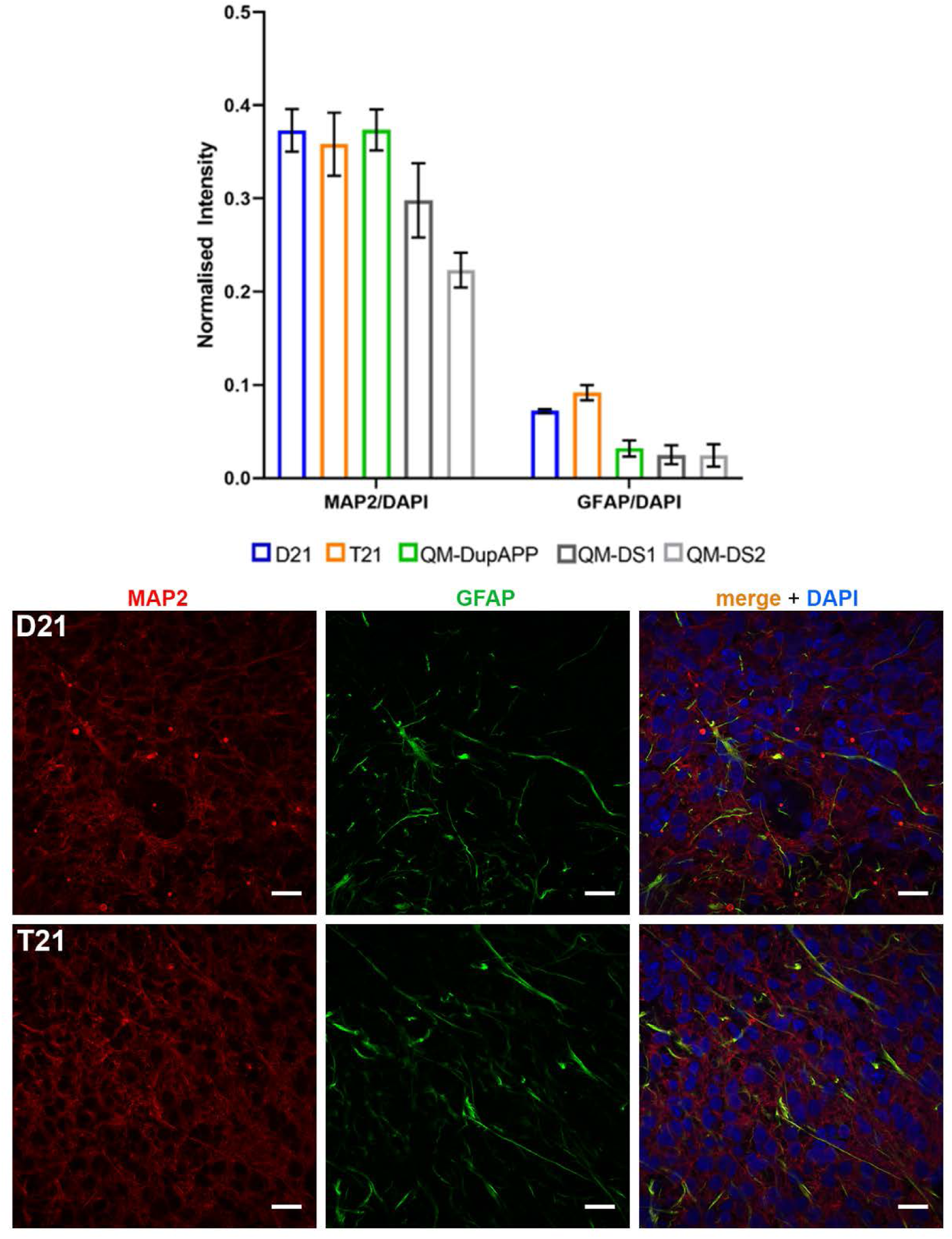
Comparison of the proportions of neurons and astrocytes to total cells in cerebral organoids. Isogenic D21 and T21 cerebral organoids, as well as organoids from DupAPP, QM-DS1 and QM-DS2 iPSCs generated mostly neurons and a small proportion of astrocytes, with none or minor differences in the proportion of astrocytes or neurons between the lines. Quantification was performed on 8 representative (Z-stack) figures each from three individual organoids per genotype. Neurons are labelled with MAP2 and astrocytes with GFAP, each marker was normalized to DAPI. Image analysis was performed using IMARIS software. Error bars: standard error. Scale bar: 20 μm.

**Supplementary Fig. 3.**
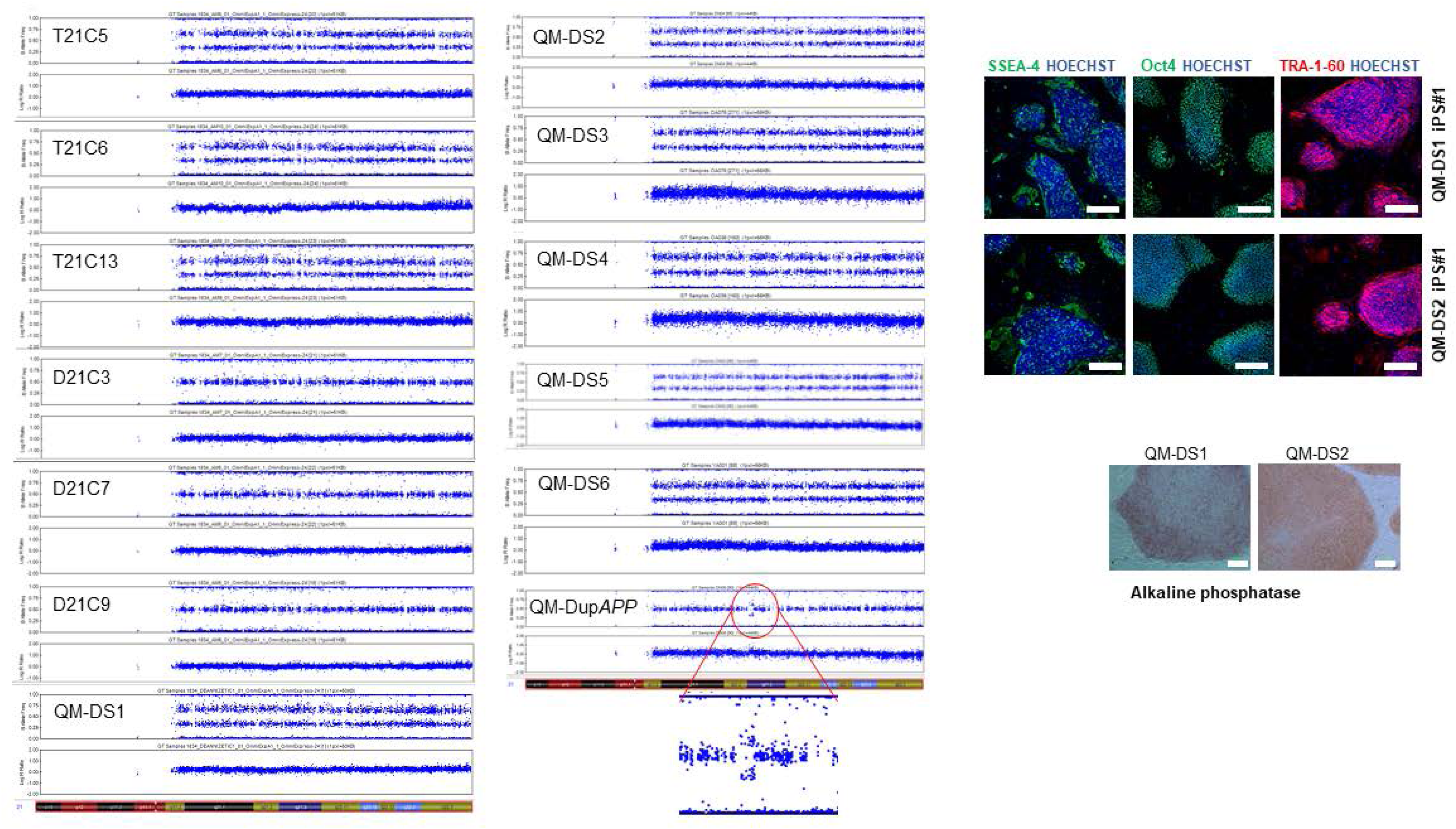
SNP arrays confirmed trisomy of chromosome 21 in all the iPSC lines used in this study, or in the case of QM-DupAPP, the partial duplication of a 580kbp segment of chromosome 21. The duplicated region in QM-DupAPP is also shown in a magnified image. Representative images of QM-DS1 and QM-DS2 confirm the expression of pluripotency markers (SSEA-4, Oct4 and Tra 1-60), scale bar: 300μm, and Alkaline Phosphatase activity, scale bar: 100μm.

**Supplementary Fig. 4.**
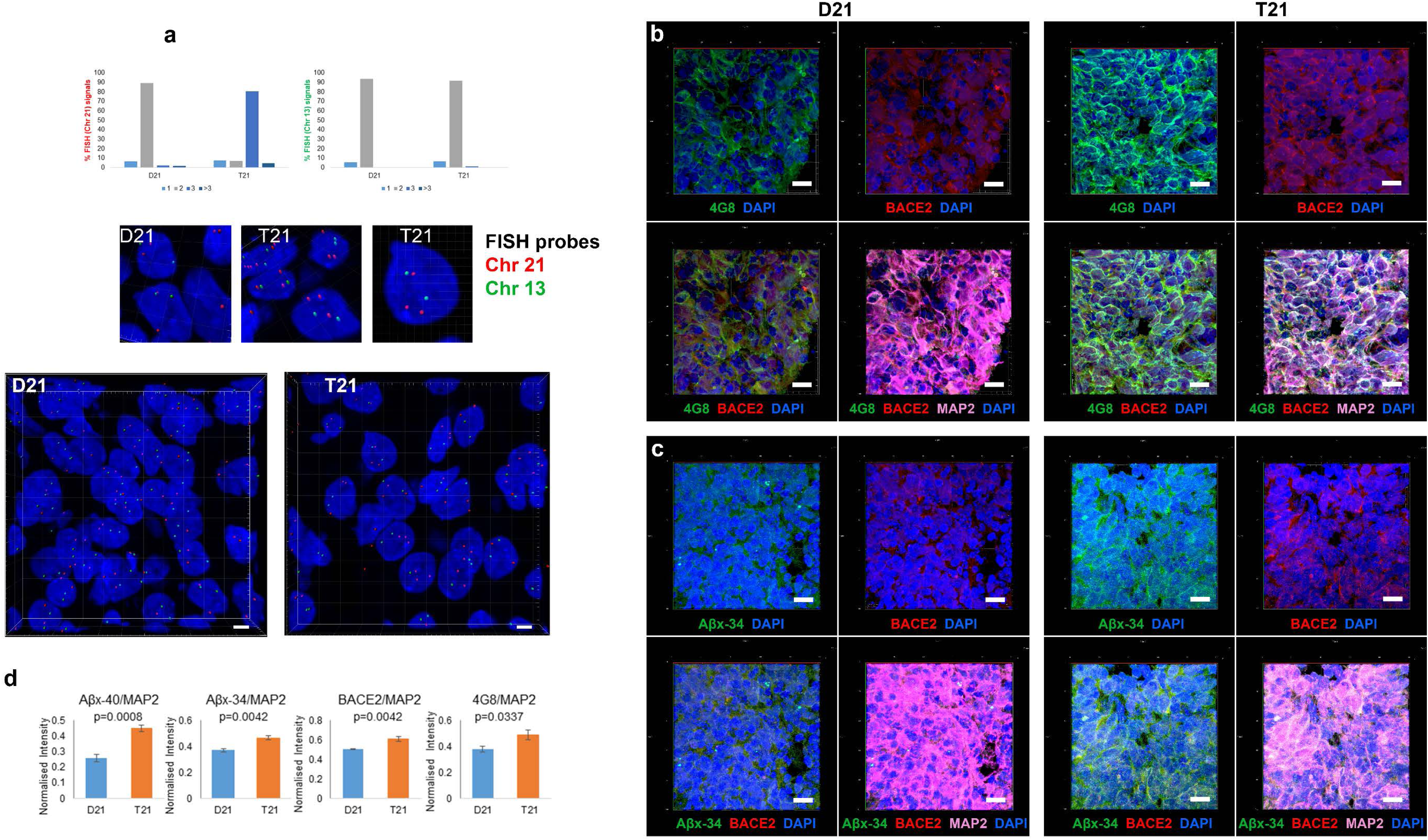
Quantitative comparison of 100DIV isogenic T21 and D21 organoid “cortical” regions by FISH and I.F. **a** For Fluorescence In Situ Hybridisation (FISH), organoid slices were processed and probed with a MetaSystems Probes for chromosome 21 (red) and chromosome 13 (green). Quantification was performed automatically using IMARIS software (nuclei were scored for 1, 2, 3 or >3 spots n>500 nuclei for each line). Scale bar: 3μm. **b** APP-4G8 antibody (green), BACE2 (red), MAP2 (magenta), DAPI (blue). Scale bar: 10μm. **c** Aβx-34 neo-epitope specific antibody (green), BACE2 (red), MAP2 (magenta), DAPI (blue). Scale bar: 10μm. **d** Quantification was performed blinded to the genotype, on 5 independent images representing three individual organoids per genotype, and containing 3,000-4,000 cells per image. Only images within the “cortical” part of the organoid were considered for the analysis. Graphs show total fluorescence intensity of positive signals for each wavelength for a given antibody, normalised by the total fluorescence intensity of MAP2 as a pan-neuronal marker. Additional staining (not shown) was analysed for Aβx-40 neo-epitope specific antibody. Image analysis was performed using IMARIS software. Error bars: standard error, and p-values: calculated after Holm-Bonferroni correction (α=0.05) of sequential two-tailed student t-test comparisons.

**Supplementary Fig. 5.**
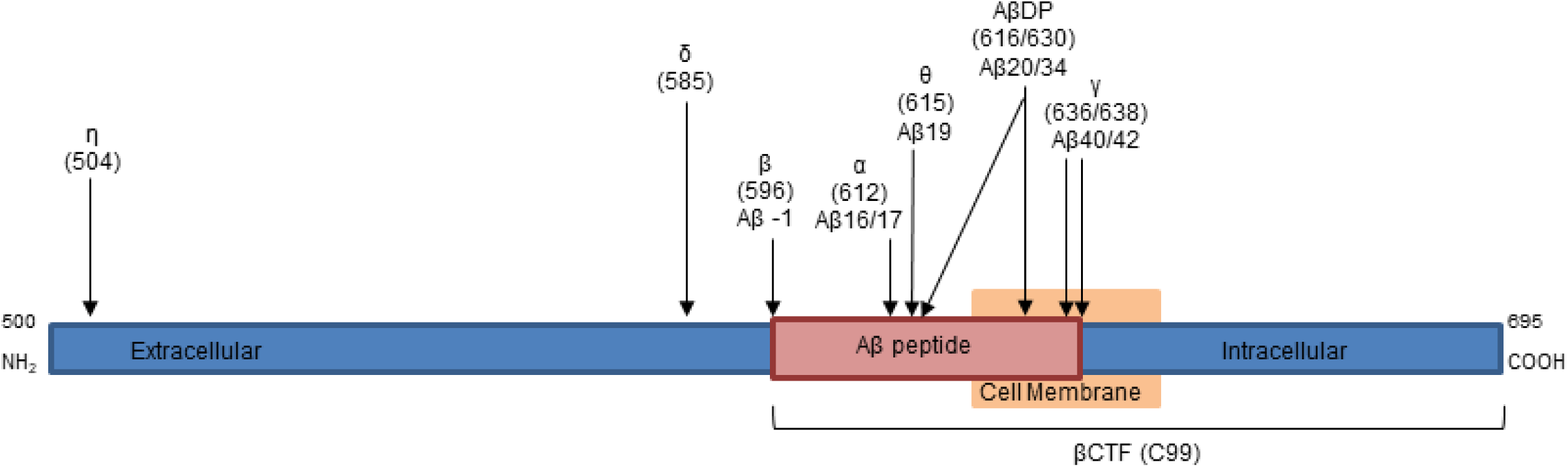
Simplified schematic representation of APP695 from amino acid 500 to the C-terminal. The amino acid cleavage site for each secretase is noted in parentheses, and for sites in Aβ the respective peptide position is also noted. Secretases for each cleavage site are η: MT5-MMP, δ: AEP, β: BACE1, α: ADAM10, θ: BACE2, γ: gamma secretase complex. In addition to it’s role as a θ-secretase, in which C99 is cleaved to preclude Aβ formation, in this report we highlight the role of BACE2 to act on Aβ as substrate, as an Aβ-degrading protease (AβDP).

**Supplementary Fig. 6.**
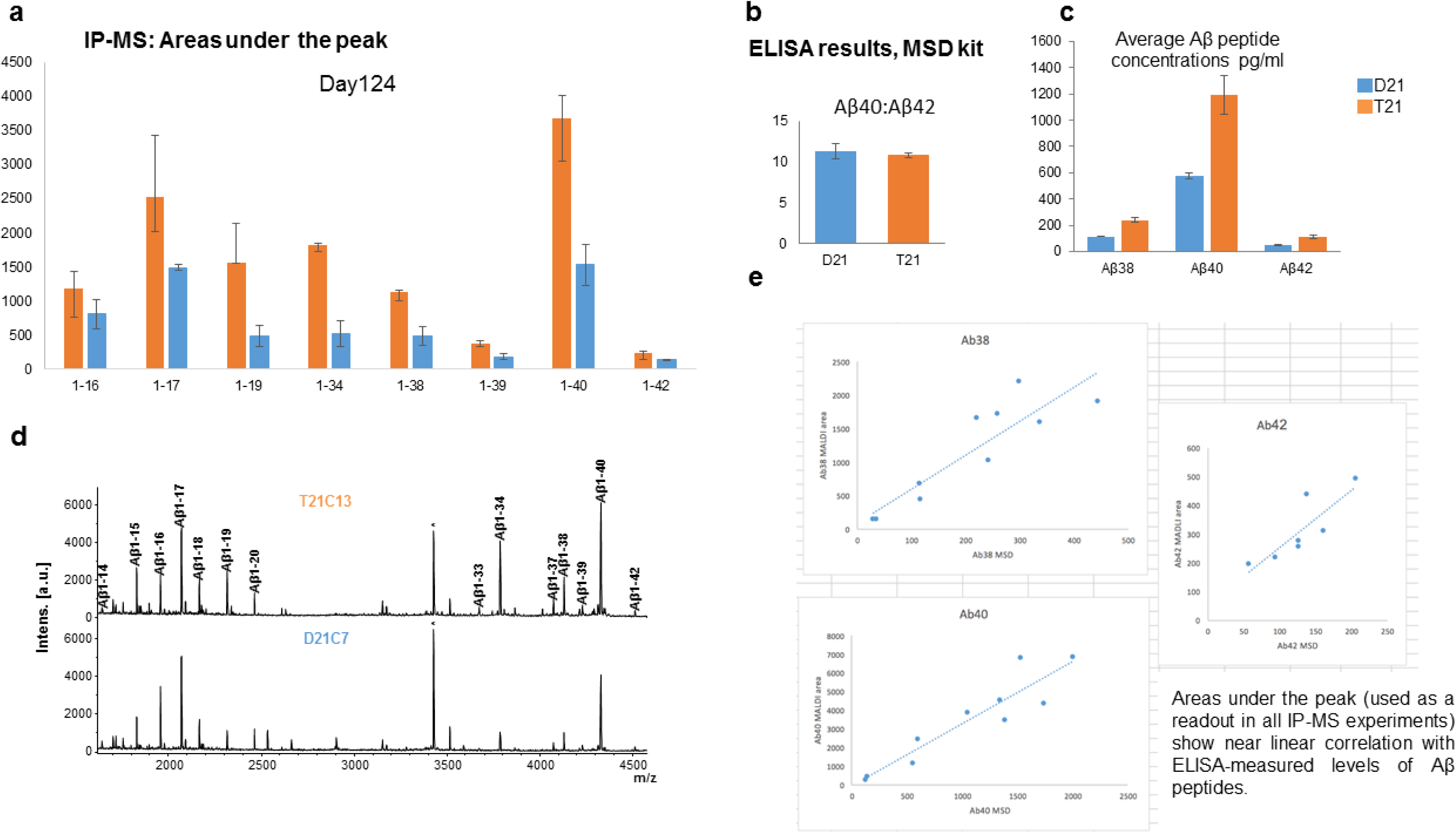
Conditioned media from isogenic D21 and T21 organoids was compared by IP-MS and ELISA. **a** IP-MS showed that all amyloid peptide species were increased in T21 compared to D21 samples. **b** Despite the increase in peptides from T21 organoids, no significant change in ratio of Aβ40:Aβ42 was detected by ELISA **c** The increase in peptide concentrations for Aβ38, Aβ40 and Aβ42 in T21 organoid CM was confirmed by MSD ELISA. **d** representative Aβ IP-MS spectra of organoid CM (*co-IP-ed unknown non-β- amyloid peptide). **e** Areas under the peak (used as a readout in all IP-MS experiments) show near linear correlation with ELISA-measured levels of Aβ peptides.

**Supplementary Fig. 7.**
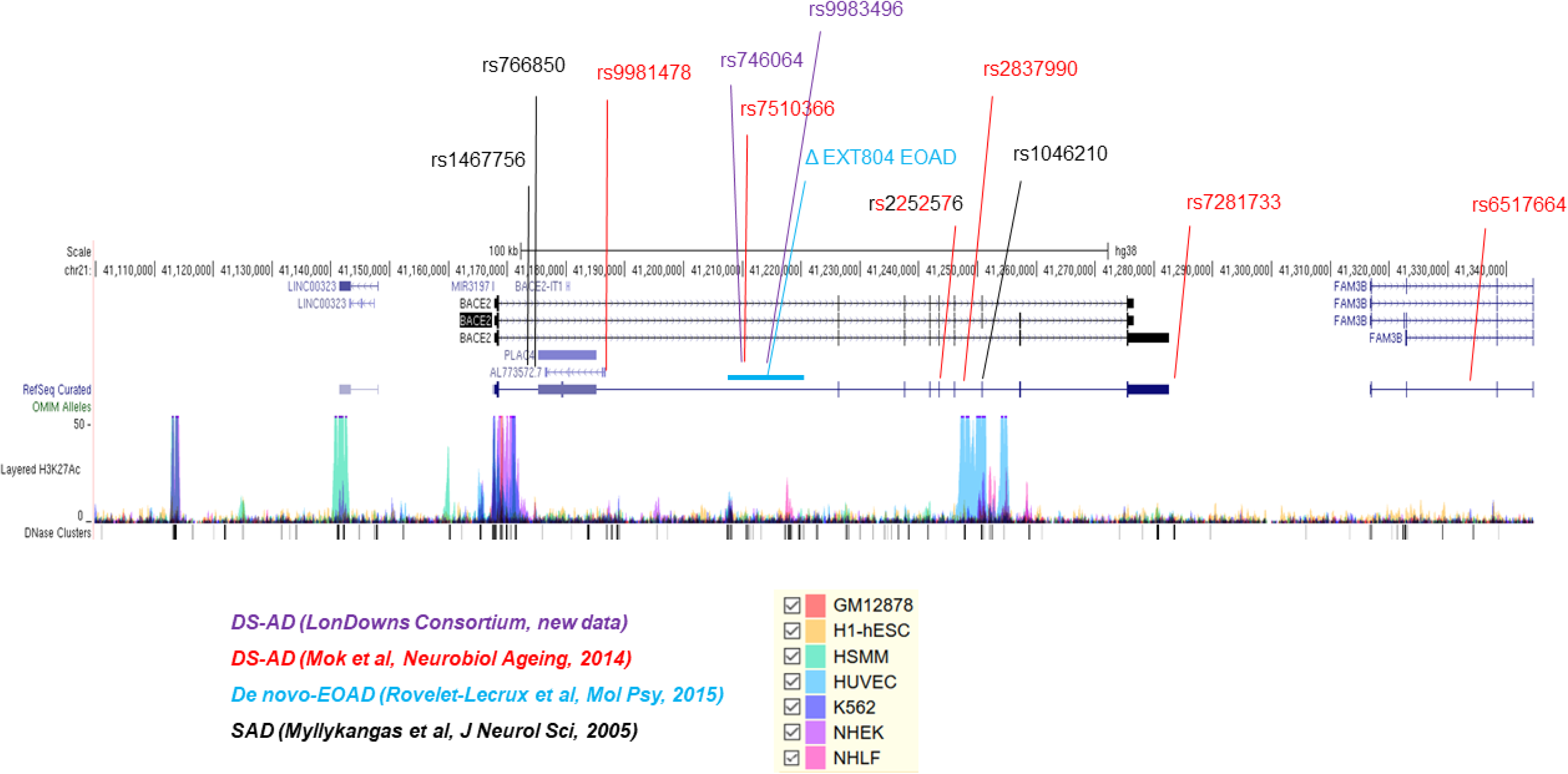
Two new single nucleotide polymorphisms in BACE2 intron1 correlate with age-of-dementia-onset among individuals with DS, and co-localize with a de-novo deletion causing non-DS EOAD. Note: two new SNPs (rs746064 and rs9983496 new data in this paper) correlating (with nominal statistical significance) with the age of dementia onset found by the genetic analysis of the LonDownS cohort of people with DS (n=554) (purple font) cluster in close proximity with one previously published SNP (red font), and all 3 SNPs are contained within a 12 kb intron1 deletion (blue line) that on its own causes EOAD in a non-DS patient. Genomic position of SNPs which have been implicated previously in sporadic Alzheimer’s disease onset in Finnish population (black font). UCSC browser snapshot of human Chr21 region 41,100,000-41,345,000 (GRCh38/hg38) containing the BACE2 locus using Gencode v24 and curated RefSeq transcript annotation is given. Underneath, the layered H3K27ac track (ENCODE) indicates putative promoter and enhancer elements present in a panel human cell lines (see legend on the right). DNase clusters (ENCODE) mark accessible chromatin.

**Supplementary Fig. 8.**
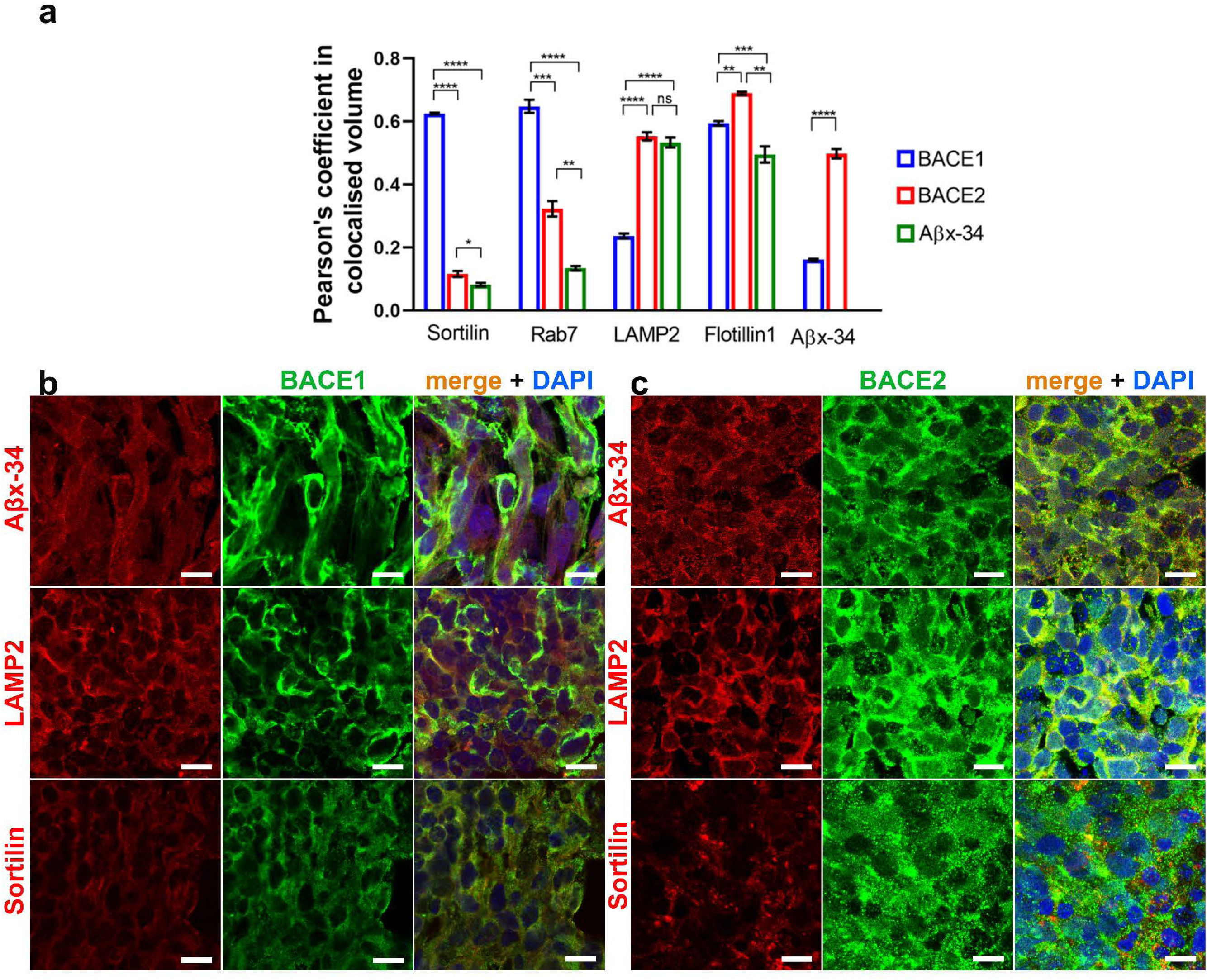
Aβx-34 colocalises with BACE2 much more than with BACE1 in T21 cerebral organoids. **a** Pairwise Pearson’s coefficient of colocalised volume for a pair of co-stained antibodies: Aβx-34 in all combinations, either BACE1 or BACE2 as a second antibody, and a marker of the sub-cellular vesicle compartment (shown at the bottom of each 3-columns histogram) as a third antibody. Error bars: standard error, and p-values: standard one-way ANOVA using post-hoc Bonferroni correction for multiple comparisons calculation. Representative individual z-slice images for the calculations performed in a are shown in **b** and c. b BACE1 (green), **c** BACE2 (green), with either Aβx-34, Sortilin or LAMP2 (red) and DAPI (blue). Scale bar: 10μm.

**Supplementary Fig. 9.**
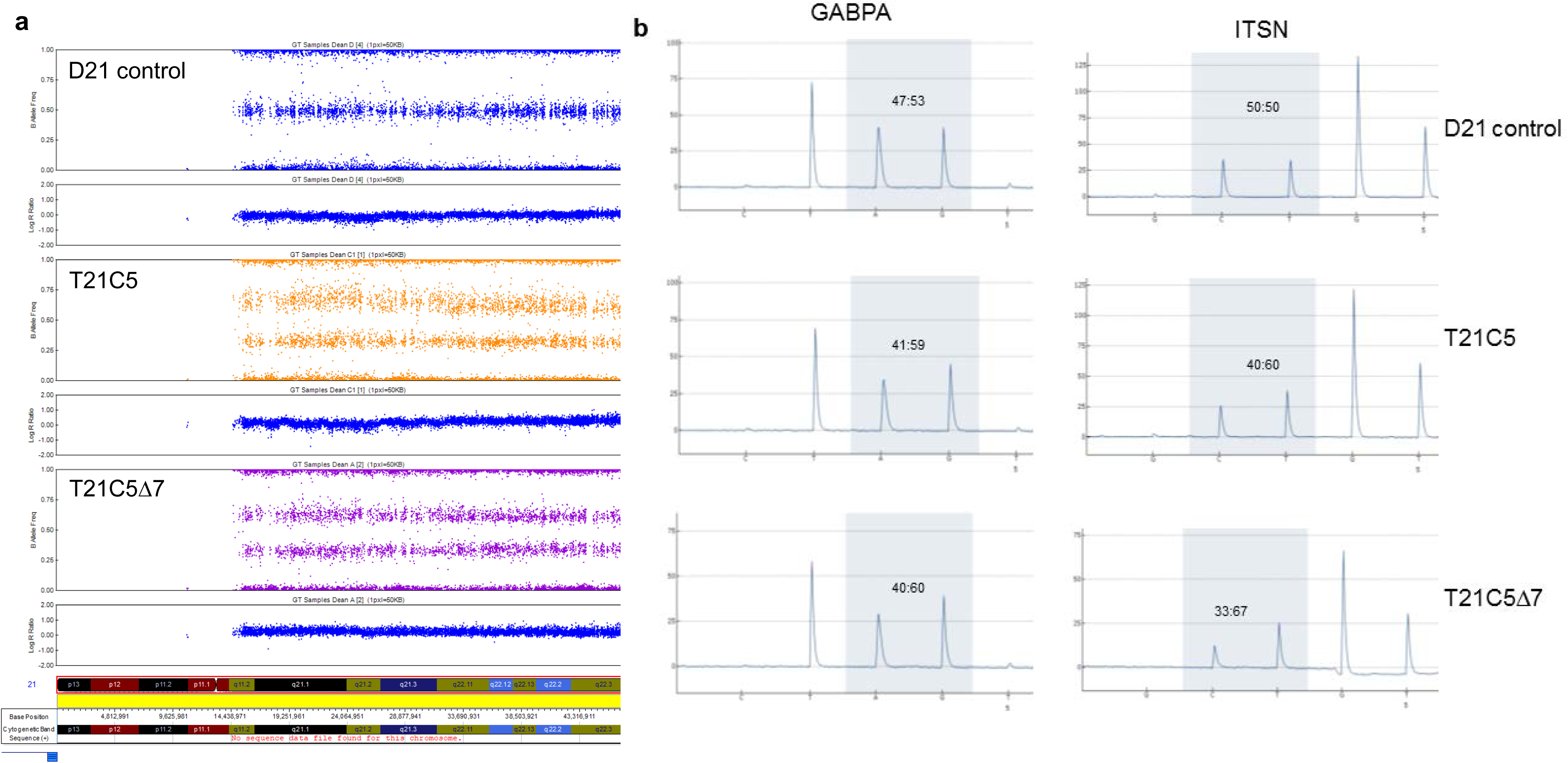
Validation of the genome integrity of CRISPR-edited iPSCs by SNP array and paralogous amplification quantitative pyrosequencing. **a** Following CRISPR editing, C5Δ7 iPSC line was assessed by SNP array and no genomic alternations were detected compared to the parental C5 iPSC line. B-allele frequency and LogR ratio plots comparing these two lines are shown for chromosome 21. Data for the whole genome available on request. **b** To further confirm retention of trisomy after BACE2 CRISPR editing, selected other genes on chromosome 21 were validated as trisomic using quantitative paralogous amplification/pyrosequencing method. GABPA and ITSN allele number were quantified relative to paralogous sequence mismatches on other chromosomes. GABPA and ITSN both show approximate 60:40 ratios for trisomic cells, and 50:50 ratios for disomic cells as expected. Quantified nucleotides are shown on the pyrogram in shaded grey boxes and the relative values for the corresponding peaks are shown.

**Supplementary Fig. 10.**
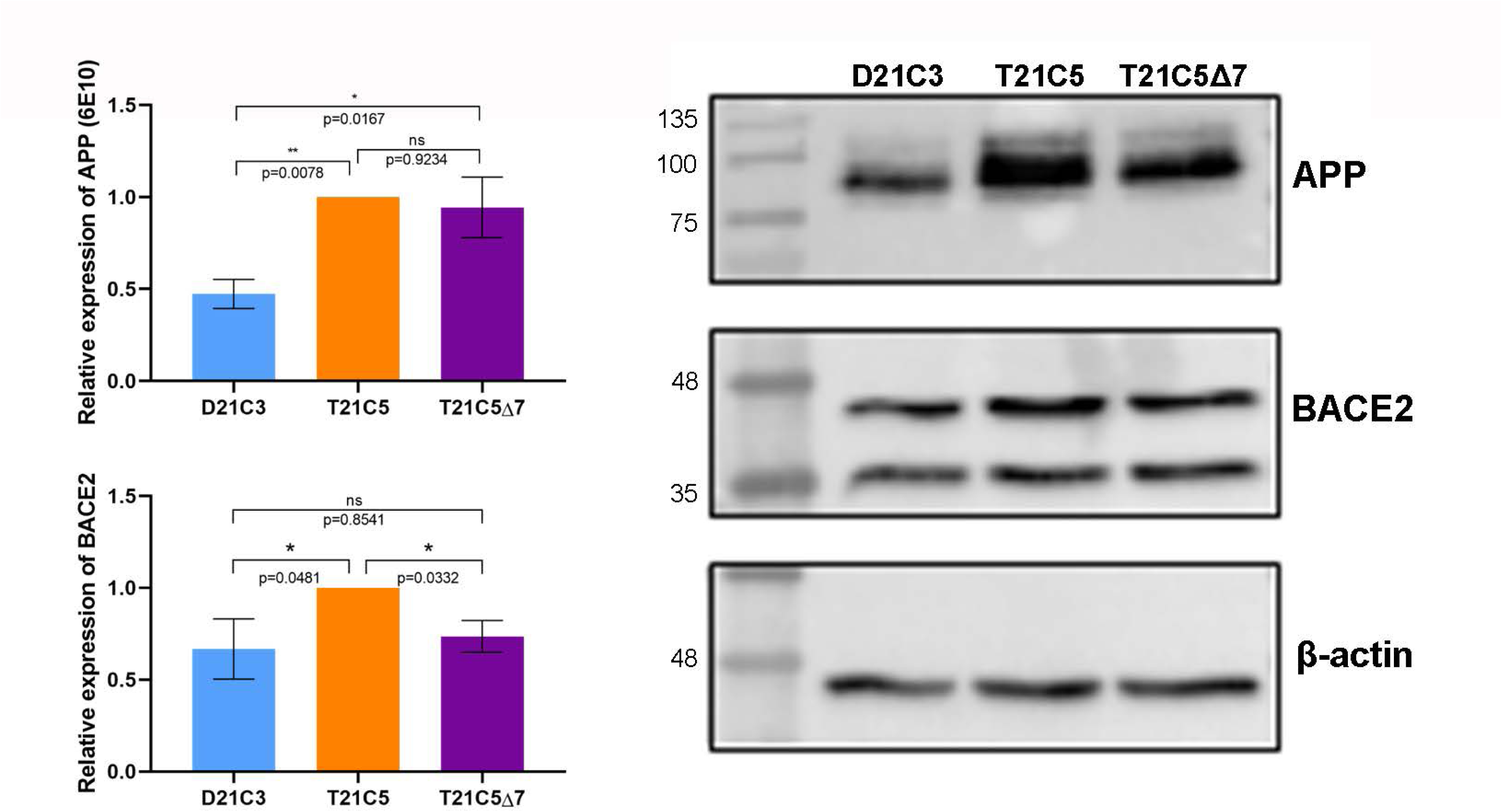
CRISPR/SpCas9-HF1-mediated reduction of BACE2 copy number from 3 to 2 in the T21C5 hiPSC line, reduced BACE2 protein expression to disomic levels, but does not alter the level of APP protein. Western blot stained with anti-BACE2 antibody or anti-APP antibody of the lysates of the iPSC line Δ7 compared to the wt T21C5, and D21C3 iPSC lines. Quantification of the total actin-normalised BACE2 signal showed a 27% reduction in Δ7 compared to T21 unedited line, and no significant difference compared to D21 control. Quantification of the total actin-normalised APP signal showed no significant difference between Δ7 and unedited T21 line, whereas they both had significantly higher APP protein levels compared to the disomic control line. Error bars: standard error, p-values after standard one way ANOVA and Tukey’s multiple comparisons test.

**Supplementary Fig. 11.**
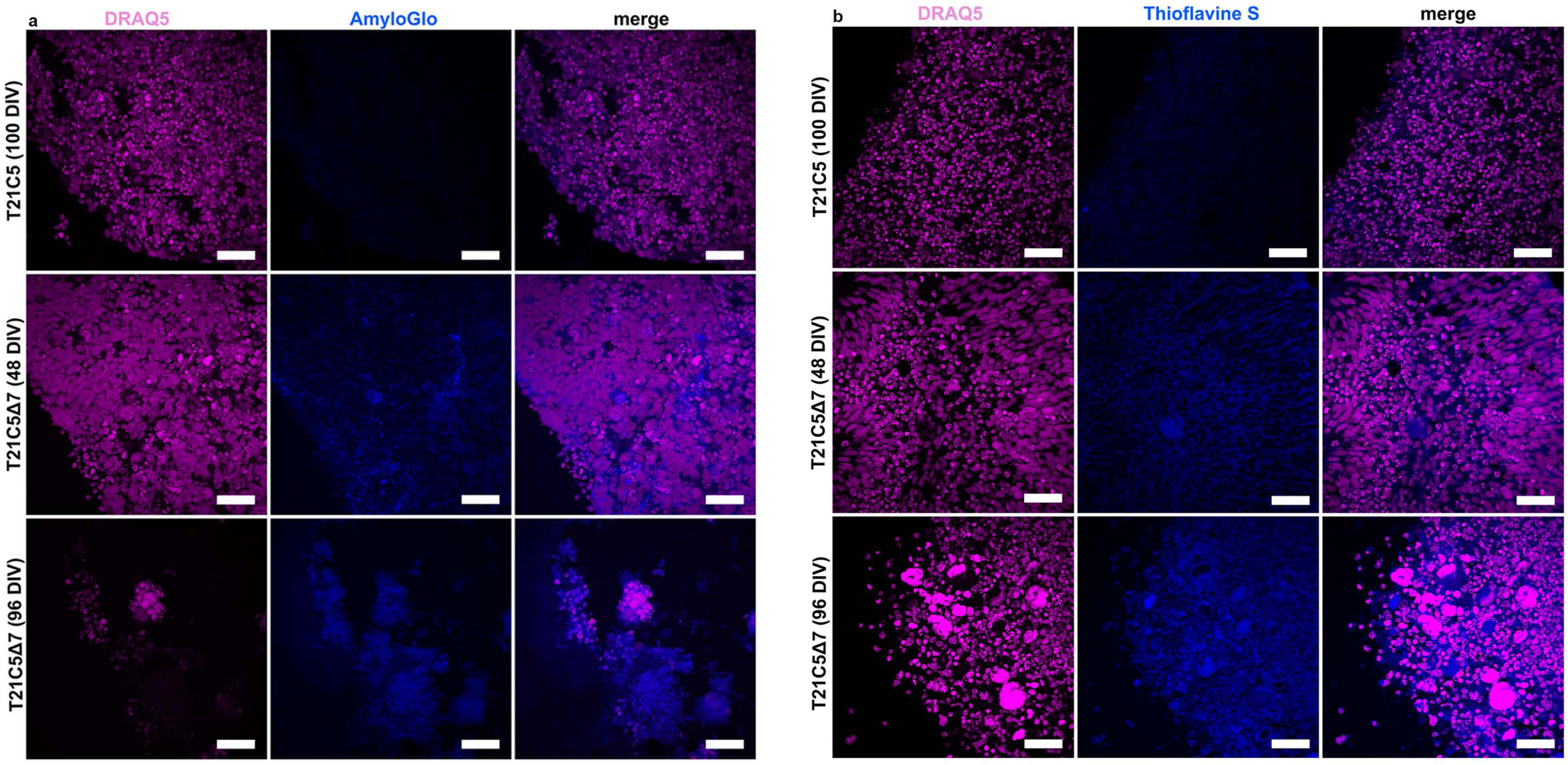
Staining of extracellular β-amyloid deposits in organoids with two different methods. **a** Early AD-like pathology was provoked in the Δ7 organoids. Staining with amyloid specific dye (AmyloGlo) and nuclear dye (DRAQ5). Top row: wt unedited control T21C5 (100DIV), Middle row: T21C5Δ7 (48DIV) and bottom row: T21C5Δ7 (96DIV); amyloid deposits are seen in Δ7 after 48DIV, but not its parental clone, at same organoid age. **b** Staining with Thioflavine S shows the same plaque-like pathology observed using AmyloGlo in the Δ7 after 48DIV. Scale bar: 20μm.

**Supplementary Fig. 12.**
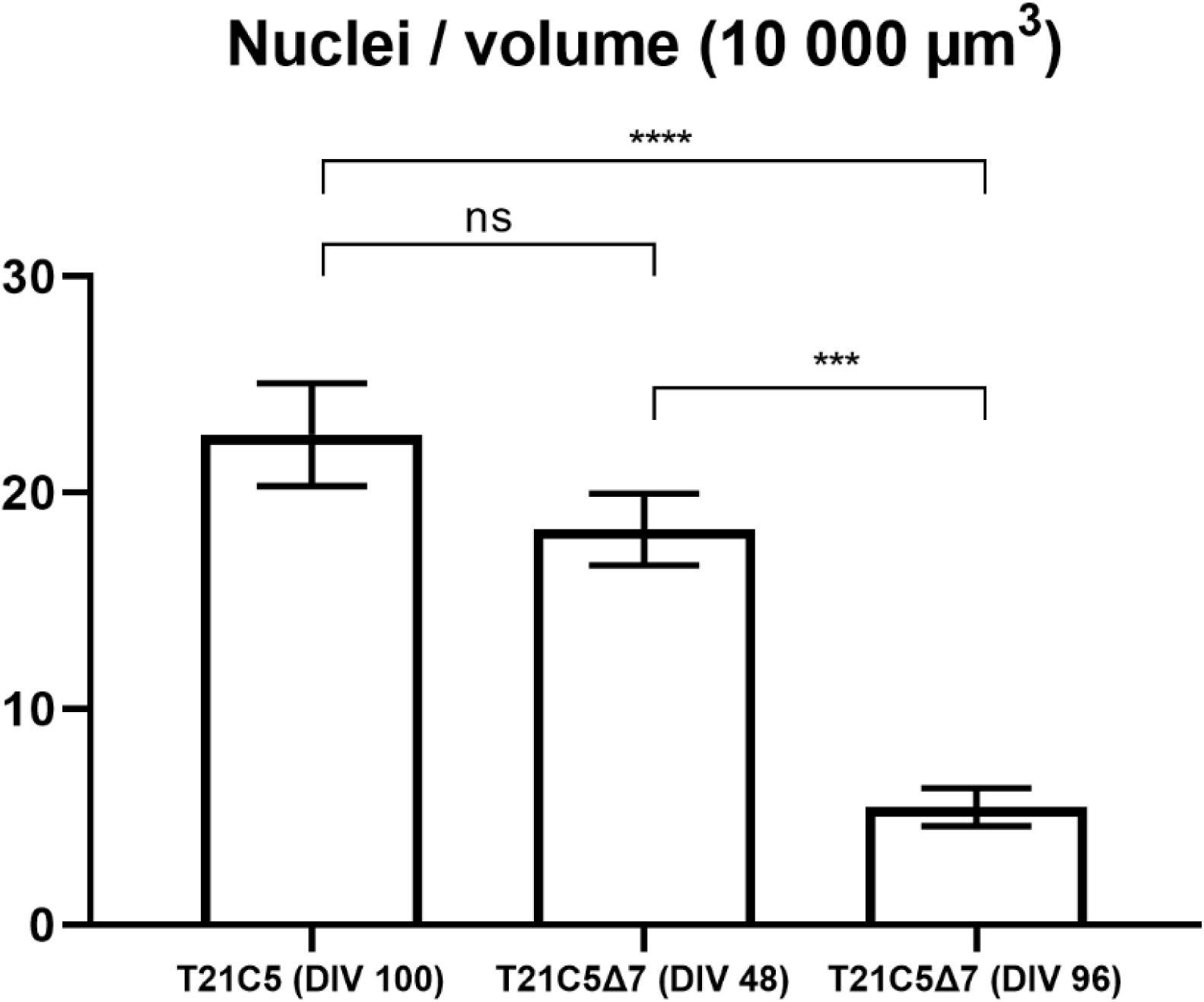
Cell death and neuronal loss in CRISPR-edited T21C5Δ7 organoids. Quantification was performed on 8 representative (Z-stack) figures from three individual organoid per genotype. Number of DAPI+ nuclei are shown in the volume of 10 000 µm^3^. Graph show decreased number of nuclei in CRISPR-edited T21C5Δ7 (DIV48) organoids compared to parental T21C5 organoids and significantly decreased number of nuclei in 96DIV organoids (p<0.0001). Significantly decreased number of nuclei were observed between DIV48 and DIV96 in CRISPR-edited T21C5Δ7 (p<0.001). Error bars: standard error, and p-values after standard one-way ANOVA and Tukey’s multiple comparison test.

**Supplementary Fig. 13.**
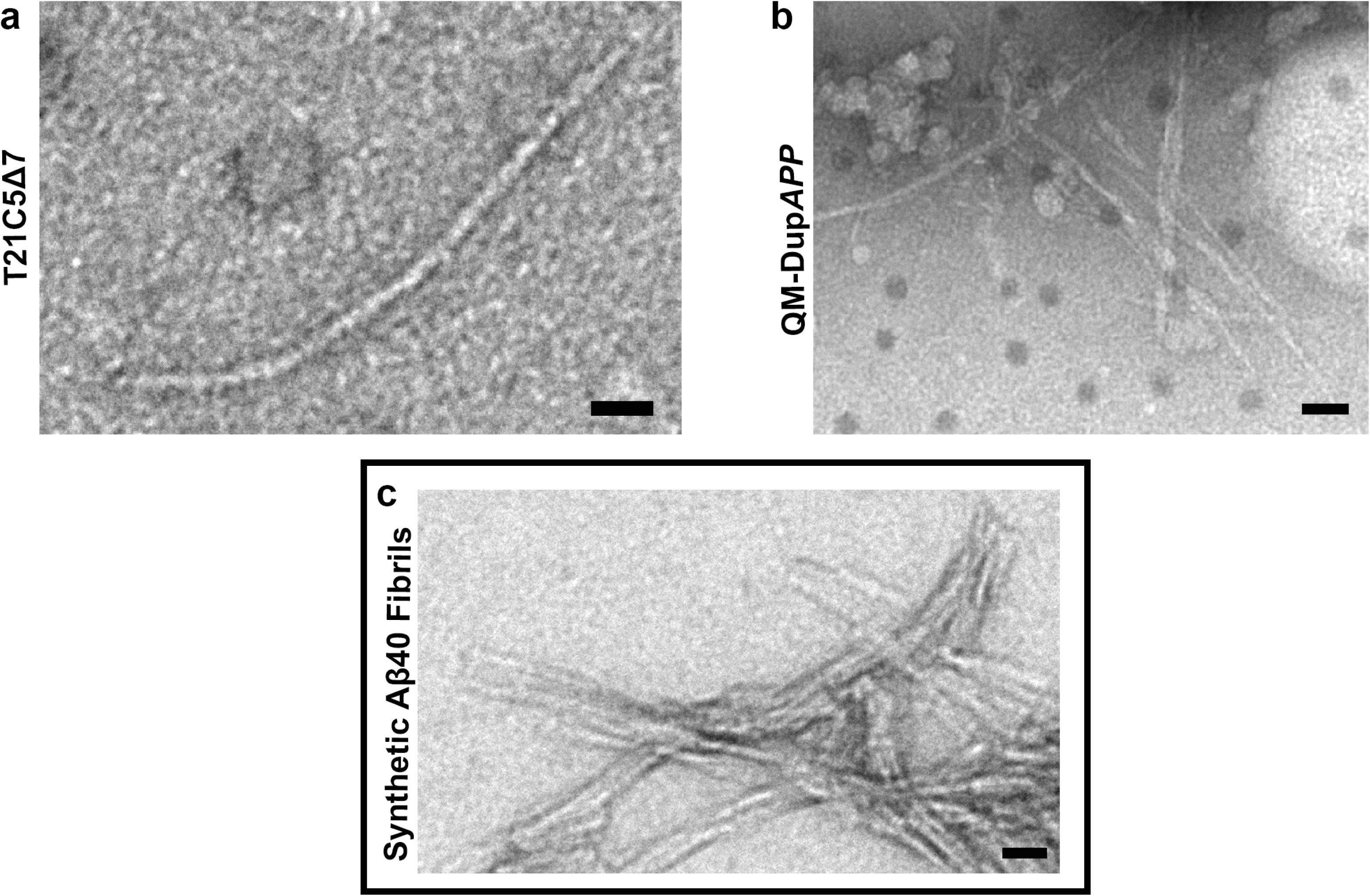
Electron micrographs of negatively stained filaments isolated from insoluble fraction of the AD-like pathology containing organoid lysates. **a, b** representative straight filaments found in the lysates from the organoids T21C5Δ7 and QM-DupAPP, respectively. **c** Aβ1-40 synthetic peptide fibrils grown in vitro. Scale bars: a, c: 20nm, b: 40nm.

**Supplementary Fig. 14.**
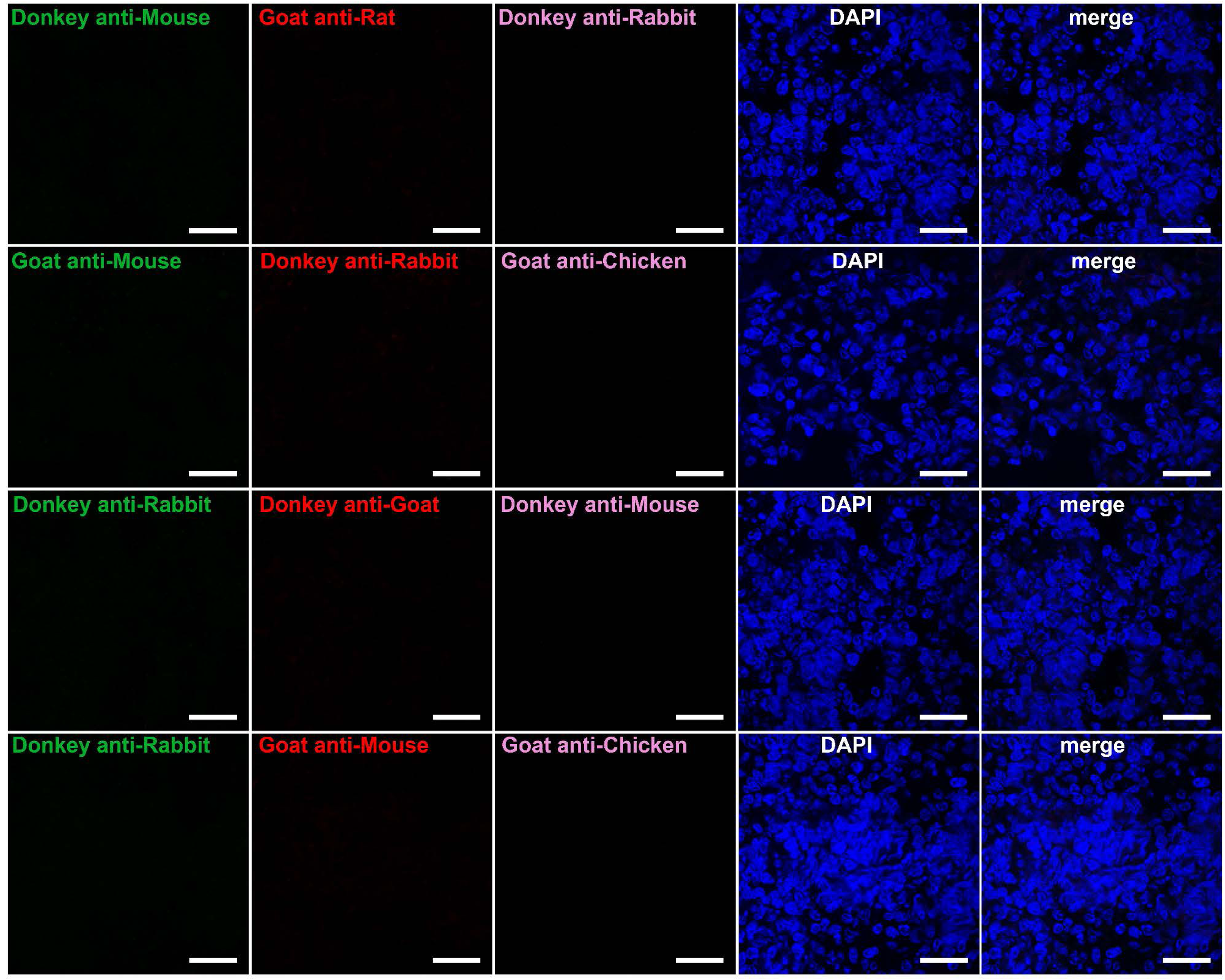
Secondary antibody alone controls for organoid immunostaining. DAPI staining confirms the presence of cells, but no unspecific signal from secondary antibodies. Scale bar: 20μm.

**Supplementary Fig. 15.**
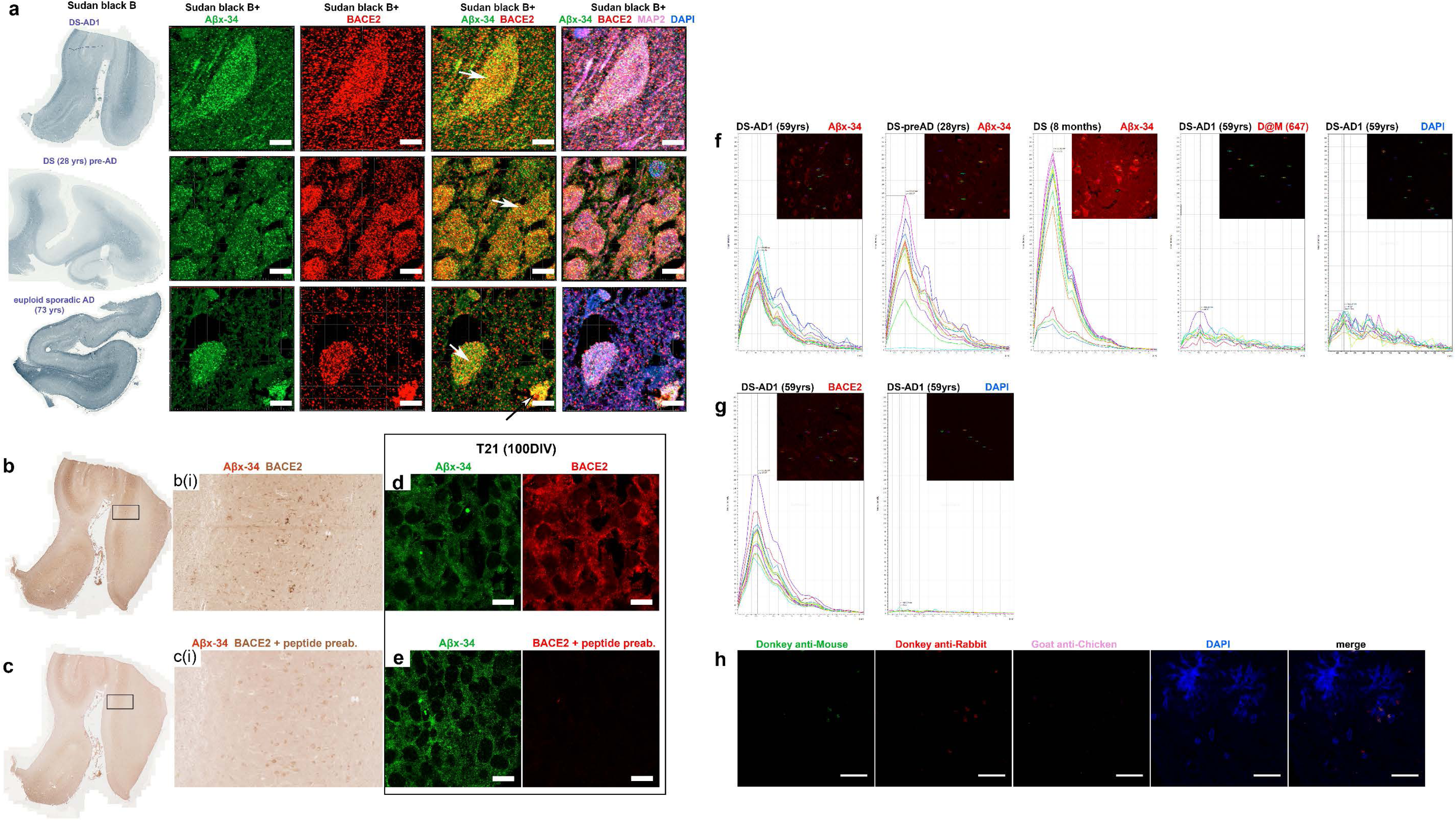
Validation and controls for immunohistochemistry. **a** Sudan black B was used to confirm the specificity of the Aβx-34 and BACE2 antibodies in human brain sections, and to eliminate lipofuscin autofluorescence. Three different human brain samples were used: DS-AD1, DS (28 yrs) pre-AD and euploid sporadic AD (73 yrs). Both antibodies show the same pattern of expression and colocalisation after Sudan black B staining (white arrows: intraneuronal fine-vesicular pattern and black arrows with white arrowhead: amorphous extra-cellular aggregates) except for a loss of the large intraneuronal spherical granules (white arrowheads, Fig. 3), which are likely lipofuscin. Scale bar: 5μm. **b** and **c** Chromogenic, immunohistochemical analysis of the human brain sections of DS-AD1, stained using polymer-HRP/AP double-staining kit. **b** The primary antibody against BACE2 was labelled with DAB (brown) and primary antibody against Aβx-34 neo-epitope was labelled with GBI-permanent-red (red); **b(i)** is a zoomed-in inset of the rectangle in B. c same as b, but both antibodies were pre-absorbed for 12 hours, and incubated overnight, with the excess of immunogenic peptide for the BACE2 antibody; **c(i)** is a zoomed-in inset of the rectangle in c. d and e: BACE2 antibody specificity control for immunofluorescence on T21 organoids (100DIV). Scale bar: 10μm. **d** Immunofluorescent staining with Aβx-34 and BACE2, **e** same as **b** and **d**, but both antibodies were pre-absorbed for 12 hours, and incubated overnight, with the excess of immunogenic peptide for the BACE2 antibody. The specificity for the neo-epitope specific antibody against Aβx-34 was extensively proven in a previous report (Cabrera et al., 2018). **f** and **g** In order to distinguish the contribution of lipofuscin auto-fluorescence to the colocalised signals, specificity of primary antibodies (Aβx-34 and BACE2) has been validated using Lambda (λ) scan function on confocal microscope (see Methods). **f** Aβx-34 shows specific peak in different ROI and uniform pattern on the three different human brain samples: DS-AD1 (59 yrs), DS (28 yrs) pre-AD and DS (8 months). As negative control of staining, DAPI and secondary antibody alone were used. **g** BACE2 also shows specific peak in different ROI and uniform pattern in human brain. **h** secondary antibody alone control. Scale bar: 20μm.

**Supplementary Table 1.**
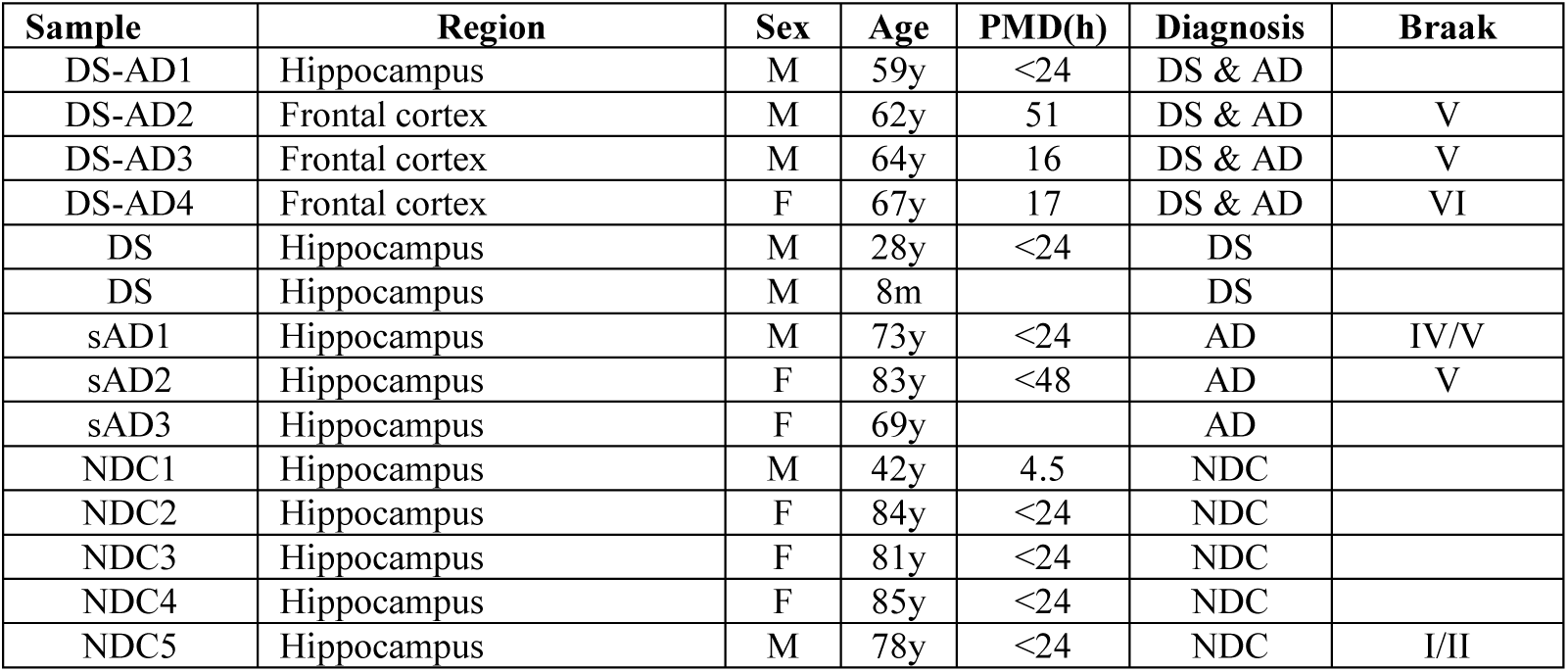
Human Brain Samples:

**Supplementary Table 2.** Chemicals:

| Chemical | Source | Cat.No | Dilution/final concentration |
| --- | --- | --- | --- |
| AmyloGlo | Biosensis | TR-300-AG | 1:100 |
| Beta Secretase Inhibitor IV (βI-IV) | Calbiochem | 565788 | 2.5μM (organoids)<br>0nM-1μM (FRET) |
| Compound E (Gamma Secretase Inhibitor XXI) | Millipore | 565790 | 6nM |
| Dako Fluorescence Medium | DAKO | S3023 | N/A |
| DAPI | Sigma | D9542 | 1:8000 |
| DAPT (Gamma Secretase Inhibitor) | Merck | 565770 | 0nM-1μM |
| DMSO | Sigma | D2650 | 1:117 |
| DRAQ5 | Abcam | ab108410 | 1:1000 |
| Formic acid | Sigma | F0507 | 87% |
| Hoechst 33342 | Life Technologies | R37605 | 1:1000 |
| LY2886721 (Beta secretase inhibitor) | Selleckchem | S2156 | 0nM-0.5μM |
| Sudan black B | Abcam | ab146284 | 0.1% in 70% EtOH |
| Thioflavine S | Sigma | T1892 | 1:100 |

**Supplementary Table 3.** Primary antibodies:

| antibody | clone | species | source | Cat.No | dilution |  |  |
| --- | --- | --- | --- | --- | --- | --- | --- |
|  |  |  |  |  | Organoids | Brain | WB |
| Aβx-34 | 1B5.4 | mouse | Ref: *Cabrera E et al. 2018* | NA | 1:1000 | 1:500 | |
| Aβx-40 | BA27 | mouse IgG2a | Wako | 014-26923 | 1:500 | 1:250 | |
| Aβx-42 (43) | BC05 | mouse IgG1 | Wako | 010-26903 | 1:500 | 1:250 | |
| Aβ | 4G8 | mouse IgG2b | BioLegend | 800701 | 1:200 | 1:200 | |
| Aβ-pE3 peptide | D5N5H | rabbit IgG | Cell Signaling | #14975 | 1:200 | | |
| APP | 6E10 | Mouse IgG1 | Biolegend | 803001 |  |  | 1:1000 |
| BACE1 |  | rabbit IgG | Abcam | ab2077 | 1:200 | 1:200 |  |
| BACE2 |  | rabbit IgG | Abcam | ab5670 | 1:200 | 1:200 | 1:500 |
| BACE2 |  | rabbit IgG | Abcam | ab5671 | 1:200 | 1:200 | 1:500 |
| BACE2 |  | rabbit IgG | Abcam | ab8025 | 1:200 | 1:200 |  |
| Beta-Actin |  | rabbit IgG | Abcam | ab8227 |  |  | 1:10000 |
| BRN2 |  | goat IgG | Santa Cruz | SC-6029 | 1:250 |  |  |
| CTIP2 | 25B6 | rat IgG2a | Abcam | ab18465 | 1:100 |  |  |
| EEA1 | C45B10 | rabbit IgG | Cell Signaling | #3288 | 1:200 |  |  |
| Flotillin1 |  | goat IgG | Abcam | ab13493 | 1:200 |  |  |
| FOXG1 |  | rabbit IgG | Abcam | ab18259 | 1:400 |  |  |
| GFAP |  | chicken IgY | Abcam | ab4674 | - | 1:250 |  |
| GFAP | 2.2B10 | rat IgG2a | ThermoFisher Scientific | 13-0300 | 1:1000 | - |  |
| HSC70 | 1B5 | rat IgG2a | Abcam | ab19136 | 1:200 |  |  |
| LAMP1 | D2D11 | rabbit IgG | Cell Signaling | #9091 | 1:200 |  |  |
| LAMP2 | GL2A7 | rat IgG | ThermoFisher Scientific | MA1-165 | 1:100 | 1:100 |  |
| LAMP2A |  | rabbit IgG | Abcam | ab18528 | 1:200 |  |  |
| LC3A | D50G8 | rabbit IgG | Cell Signaling | #4599 | 1:400 |  |  |
| MAP2 |  | chicken IgY | Abcam | ab5392 | 1:1000 | 1:500 |  |
| Rab7 | EPR7589 | rabbit IgG | Abcam | ab137029 | 1:200 |  |  |
| Rab7 | Rab7-117 | mouse IgG2b | Abcam | ab50533 | 1:200 |  |  |
| REELIN | 142 | mouse IgG1κ | Chemicon (Merck) | mab5366 | 1:300 | | |
| SATB2 |  | IgG | Abcam | ab34735 | 1:200 |  |  |
| Sortilin |  | goat IgG | R&D Systems | AF3154 | 1:200 |  |  |
| Tau (3-repeat isoform RD3) | 8E6/C11 | mouse | Millipore | #05-803 | 1:500 |  | 1:1000 |
| Tau (hyperphosphorylated) | AT8 | mouse IgG | ThermoFisher Scientific | MN1020 | 1:500 |  |  |
| Tau (filamentous) | AT100 | mouse IgG | ThermoFisher Scientific | MN1060 | 1:1000 |  |  |
| Tau (conformationally altered) | TG3 | mouse IgG | From Peter Davies (via Alzforum) | NA | 1:100 |  | 1:1000 |
| TBR1 |  | rabbit IgG | Abcam | ab31490 | 1:500 |  |  |

**Supplementary Table 4.** Secondary antibodies:

| antibody | conjugate | source | Cat.No | dilution |
| --- | --- | --- | --- | --- |
| Donkey anti-Mouse IgG (H + L) | Alexa Fluor 488 | ThermoFisher Scientific | A-21202 | 1:1000 |
| Donkey anti-Mouse IgG (H + L) | Alexa Fluor 555 | ThermoFisher Scientific | A-31570 | 1:1000 |
| Donkey anti-Mouse IgG (H + L) | Alexa Fluor 647 | ThermoFisher Scientific | A-31571 | 1:500 |
| Donkey anti-Rabbit IgG (H + L) | Alexa Fluor 488 | ThermoFisher Scientific | A-21206 | 1:1000 |
| Donkey anti-Rabbit IgG (H + L) | Alexa Fluor 555 | ThermoFisher Scientific | A-31572 | 1:1000 |
| Donkey anti-Rabbit IgG (H + L) | Alexa Fluor 647 | ThermoFisher Scientific | A-31573 | 1:500 |
| Donkey anti-Goat IgG (H + L) | Alexa Fluor 555 | ThermoFisher Scientific | A-21432 | 1:1000 |
| Goat anti-Mouse IgG2b (H + L) | Alexa Fluor 488 | ThermoFisher Scientific | A-21141 | 1:500 |
| Goat anti-Mouse IgM (H + L) | Alexa Fluor 568 | ThermoFisher Scientific | A-21043 | 1:500 |
| Goat anti-Rat IgG (H + L) | Alexa Fluor 568 | ThermoFisher Scientific | A-11077 | 1:1000 |
| Goat anti-Chicken IgY (H + L) | Alexa Fluor 633 | ThermoFisher Scientific | A-21103 | 1:500 |
| Goat anti-Rabbit IgG (H + L) | HRP | Abcam | ab97051 | 1:10000 |
| Goat anti-Mouse IgG (H + L) | HRP | Abcam | Ab97023 | 1:10000 |
| VECTASTAIN ABC HRP Kit | HRP | Vector | PK-4002 | 1:200 |
| Double staining Kit | HRP & AP | GBI Labs | DS202A-18 | NA |

**Supplementary Table 5.**
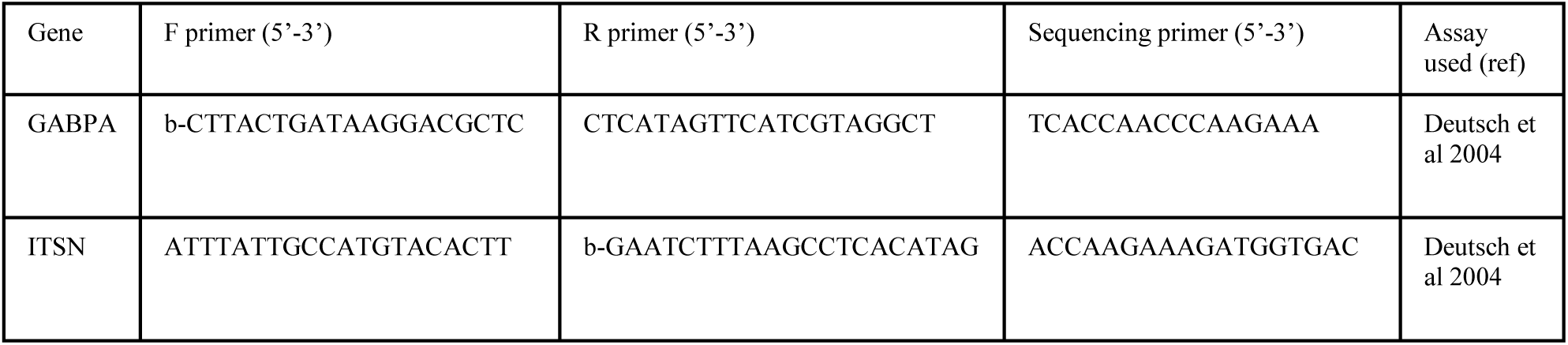
Primer sequences used for the paralogous-loci-amplification-quantitative pyrosequencing

## Supplementary Information

### Related to Fig. 1

Fig. 1a: Variability between individual iPSC lines (representing individual re-programming events) was tested by ANOVA in Exp1, where all 3 independent trisomic lines of our isogenic model were used in a single experiment. No significant differences between individual lines were found in any of the calculations shown in Fig. 1, demonstrating that our peptide-ratio-readout parameter is driven by the genotype, and not re-programming artefacts or culture history of the iPSC lines (data did not fit the allowed space, available on request).

As peptide-ratio readouts differed slightly between three independent experiments, we are showing complete data here for each experiment individually. As shown in Fig. 1a, the difference (or the absence of difference) caused by T21 in an isogenic comparison remained stable in each of 3 experiments. In Exp3, for the ratio of 1-19/amyloidogenics, the isogenic comparison of T21 v D21 showed a p=0.027 (2-tailed t-test), which dropped to p=0.0681 after ANOVA comparison with all 5 individual samples.

Also in Exp3, we further performed an analysis by genotype groups. For the AβDP/amyloidogenics ratio, the combined T21 samples (n=3) were significantly higher than D21 (ANOVA p=0.0021), and significantly higher than Dup*APP* (ANOVA p=0.0011), whereas D21 is not significantly different from Dup*APP*. The same result was obtained for the total BACE2/amyloidogenics ratio: combined T21 (n=3) v D21, ANOVA p=0.0138; combined T21 (n=3) v Dup*APP*, ANOVA p=0.0036, and D21 v Dup*APP* shows no significant difference. The comparison of α-site cleavages (1-16&1-17)/amyloidogenics never showed any significant difference irrespective of how the samples were grouped.

### Related to Fig. 2

Fig. 2: The FRET assay positive control was performed using recombinant human BACE2 at 37⁰C, pH=3.5 for 2h in the R&D systems assay buffer, as specified in the manufacturer’s protocol, using the R&D systems FRET control peptide (ES010). In three technical replicates the blank-subtracted raw fluorescence readings obtained were 13,836(±130 SEM). BACE2 with the new FRET peptide for the AβDP cleavage after aa34 (in the absence of any inhibitors) gave blank-subtracted readings 10,100(±59 SEM). This was taken as the 100% value for the graphs shown in Fig. 2. For comparison, BACE1 incubated with the same FRET peptide, using the manufacturer’s assay buffer for BACE1, gave the readings of 522 (±58 SEM) in the same experiment.

### Related to Fig. 3 and Supplementary Fig. 8

We compared the degree of colocalisation between either BACE1 or BACE2, and Aβx-34 clearance product in organoids, along with other markers of intra-neuronal compartments: Flotillin1 (general marker of lipid rafts), Rab7 (late endosome marker), Sortilin (a major ApoE receptor linked to Aβ catabolism), and LAMP2 (one of the lysosomal membrane proteins often used to visualize lysosomes in studies of Aβ-processing).

Both BACE1 and BACE2, as well as Aβx-34 highly colocalised with Flotillin1, suggesting that this type of Aβ degradation takes place in lipid raft containing vesicles (Fig. 3 and Supplementary Fig. 8). However, BACE1 and BACE2 differed in vesicular sub-compartment distribution: BACE1 was highly colocalised (>0.6) with each Sortilin and Rab7 and only weakly with LAMP2 (0.22), whereas BACE2 did not co-localise with Sortilin(<0.1), but colocalised moderately with Rab7 (0.31) and highly with LAMP2 (>0.5) (Supplementary Fig. 8). Interestingly, the localisation of the Aβx-34 fragment closely resembles the pattern of BACE2, and not of BACE1: (Pearson coefficient of 0.1 with each Sortilin and Rab7, and >0.5 with LAMP2), further supporting the observation of Aβx-34 (>0.5) localisation with BACE2 and less so with BACE1, in both organoids (Fig. 3 and Supplementary Fig. 8) and human brain (Fig. 4). In order to define the compartment with the highest concentration of Aβx-34 within the endo-lysosomal system more precisely, we co-stained the Aβx-34 neo-epitope-specific antibody with other markers associated with Aβ processing: LC3A (macro-autophagosome marker), EEA1 (early endosome marker) and LAMP1 (a classical lysosome marker). Surprisingly, none of these markers showed any colocalisation, demonstrating that Aβx-34 is not present in either early endosomes, macro-autophagosomes, or classical lysosomes (Fig. 3). As Aβx-34 did not colocalise with LAMP1 or LC3A, but colocalised strongly with LAMP2, we tested a colocalisation with the components of an alternative autophagy pathway that would be compatible with this pattern of colocalisations: chaperone-mediated autophagy (CMA). Unexpectedly, we detected an extremely high level of co-localization of Aβx-34 with both HSC70 (chaperone in CMA) and LAMP2A, (the isoform of LAMP2 that is the main protein controlling the levels of CMA activity) (Fig. 3). Some intra-neuronal LAMP2A+ vesicles appear to contain both HSC70 and Aβx-34 (Fig. 3). These data suggest that AβDP activity of BACE2 is linked with the CMA pathway.

### Related to Fig. 4

Fig. 4a-d: As immunofluorescence on brain sections is susceptible to bright and false positive autofluorescent signals from lipofuscin granules, we confirmed the colocalisation of Aβx-34 and BACE2 using non-fluorescent, chromogenic dual labelled immunohistochemistry (Supplementary Fig. 15b), where the specificity of the BACE2 antibody was further verified by pre-absorption control with the immunogenic peptide (Supplementary Fig. 15c). This method confirmed the intra-neuronal co-localization of Aβx-34 and BACE2 signals.

### Related to Fig. 5

The 7bp deletion causes a frameshift at aa157 of BACE2 protein sequence. This introduces a stop codon within the protease cleavage domain at aa197. The potential off-target effects of the CRISPR guide RNA used were tested using two prediction software tools: CCTop and http://crispr.mit.edu/. No target sequences were found with 0, 1 or 2 mismatched nucleotides. No targets, that had three or more mismatches were overlapping between the two software predictions. In CCTop, only two sites with three mismatches, and more sites with four mismatches were found. Top 10 loci from this prediction were amplified with the putative target sequence in the middle, and sequenced in the T21 wt iPSC compared to the Δ7 iPSC line. No off-target effects of the CRISPR/SpCas9-HF1 intervention were detected.

